# Bioscaffold guidance drives liver periportal area tubulogenesis in hIPSC organoids

**DOI:** 10.1101/2025.08.28.672864

**Authors:** Harini Rajendiran, Geetika Sahni, Aditya Arora, Matt s. Hepburn, Flora Luciani, Ong Hui Ting, Jin Zhu, Marion Marchand, Selma Serhrouchni, Anais Monet, Piyush Mishra, Nai Mui Hoon Brenda, Jiayue li, Alireza Mowla, Farzaneh Navaeipour, Brendan f. Kennedy, Virgile Viasnoff

## Abstract

Recapitulating the liver periportal area *in vitro* remains a major challenge due to its complex cellular composition and the coordinated development of both ductal and endothelial networks. Most existing bile duct organoid models fail to reproduce tubular extension and multicellular organization. We present a 3D ECM micro-rods system biofunctionalized with grafted growth factors mimicking fetal liver paracrine signaling to initiate and to coordinate the development of bile ducts, parenchymal cells, and vascular structures. Our system recapitulates key developmental stages, from the formation of the ductal plate to the emergence of elongated tubular ducts. The ECM rod alone provides the biophysical cues initiating the elongation of the bile ducts and the spatial structuration of the cellular organization, while the biofunctionalization with growth factors enhance the organization and maturation of the biliary and vascular systems. We further demonstrate that this approach is scalable and amenable to quantitative analysis through label-free Optical Coherence Tomography combined with AI-driven 3D segmentation.

## Introduction

The development of the liver periportal area is key to the functional organogenesis of the liver lobule. In the human embryo, it occurs between EW8 and EW10 and is spatially organized around the portal vein [1, 2]. Paracrine signaling via TGF-β and Notch from adjacent endothelial and mesenchymal cells induces the differentiation of embryonic hepatoblasts into cholangiocytes in the close vicinity of the vein. In turn, cholangiocytes develop the tubular network of intrahepatic bile ducts (IHBD) that closely scaffolds the portal vein network [3–6]. At EW10, hepatic arteries begin to develop in the vicinity of the IHBD, reportedly in response to VEGF signaling from cholangiocytes [7, 8]. Meanwhile, the remaining parenchymal hepatoblasts differentiate into hepatocytes. Local paracrine signaling, acting over just a few cell diameters, is paramount to the spatial and temporal coordination of the three tubular networks (portal vein, hepatic arteries, and intrahepatic bile ducts), with each network signaling for the development and maintenance of the others.

Our current understanding of the portal triad is largely based on homologies between murine and human development. Human liver organoid models have focused either on the generation of complex liver bud surrogates that reproduce the diversity of cell types [9–11], irrespective of their spatial organization, or on the development of bile duct organoids from induced or primary cholangiocytes [12–18]. In all cases, cellular differentiation in organoids is controlled by a sequence of homogeneous soluble growth factor treatments and by embedding them in an extrinsic (culture within) extracellular matrix scaffold. Under such conditions, cholangiocytes can mature into large spherical cysts comprising a single-cell monolayer. More advanced techniques, like 2D patterning and hydrogel microparticles, have enabled spatiotemporal regulation of hepatoblast fate, but they often lack ECM-derived cues and topological drivers that are critical for cytoskeletal reorganization during tissue formation [19–21]. Whereas the osmotic growth of cholangiocytes cysts most often leads to spherical structures, recent studies using 2D patterning on fibronectin substrates have demonstrated the creation of cholangiocytes tubes based on a non-physiological self-wrapping cell layer mechanism [22]. Primary adult (and not embryonic) cholangiocytes placed in an optimized ECM environment have been shown to create branched bile duct–like tubules [23]. Lastly, the co-culture of hepatoblasts around endothelial cells adhered to a cylindrical rod led to the formation a periportal area like structure. Efforts to incorporate portal vein cues have generated bile ducts in proximity to the signaling center, but these systems remain highly work intensive, low-throughput and hardly scalable [24].

Since the liver portal triad development results principally from local juxtacrine (ie local) and paracrine (ie short range) signaling around the portal vein scaffold, we reasoned that the classical approaches in organoid culture using soluble growth factors stimulation might lack important components necessary to the portal triad development. In this paper, we used rod-like scaffolds biofunctionalized with growth factors to mimic the paracrine signaling that elicits the initial differentiation of hepatoblasts along the portal vein. In contrast to the conventional approaches requiring soluble cues and ECM embedding, our system recapitulate the periportal organogenesis from hPSCs capturing the coordinated emergence of ductal, parenchymal and vascular compartments. It reproduces a large fraction of the cellular diversity and the temporal sequence of development including the formation of a tubular bile duct. supports intrinsic self-organization of the portal triad architecture. To enable the dynamic and non-invasive morphological assessment of these structures, we developed a 3D imaging pipeline using Optical Coherence Tomography (OCT), allowing the scalable quantification of the ductal architecture and its morphogenesis. We specifically focused on the first 7 days of development during which the main morphological events occur. Tests over longer maturation periods are left for future work since they involve other classes of growth factors and micro-environment scaffolding.

## Results

### Scalable biofunctionalized 3D scaffolds mimicking portal vein cues

To recreate the spatial organization of cellular interactions present along the embryonic portal vein, we engineered a series of biofunctionalized scaffolds designed to deliver juxtacrine and paracrine cues (Notch, TGF-β, and geometry) mimicking the instructive role of the mesenchyme and endothelial lining of the vein in vivo (Figure 1a). As surrogates for the embryonic portal vein, we engineered Gelatin Methacrylate (GelMa) rods with physiological dimensions (0.3 × 1 mm) and addressable combinations of growth factors chemically conjugated to the extracellular matrix (ECM) (Figure 1b, Materials and Methods). Molding the rods in long Polyethylene tubes (*500* μm *internal diameter*) ensured consistent dimensions: diameter of 400 μm (after swelling) and lengths ranging between 0.8 to 1.5 mm after manual chopping (**Figure 1c, Supplementary** Figure 1a). The elastic modulus of the scaffolds measured using atomic force microscopy (AFM) averaged around 3.6 kPa, in agreement with the physiological range (**Supplementary figure 1b**). We coupled the growth factors (GF) to the rods using their heparin-binding domain when present. For those lacking this domain (Jagged1-FC), we employed protein G based coupling to enable targeted binding (Figure 1b). In all cases, we confirmed that the coupling was localized at the rod surface (Figure 1c). Performing ELISA on the supernatant of the incubation medium, we quantified that up to 46% of the soluble growth factors from the initial bound to the rods (**Material and Methods**) corresponding to an average of 761 pg of Activin A per rod likely saturating the biding sites. This immobilization efficiency was consistent across different ligands and did not vary with longer washing period demonstrating that growth factors are not released from the rods (**Figure 1c**, **Supplementary 1b**). We prepared the biofunctionalized rods in high-throughput batches and seeded individually into U-bottom 96-well plates to scaffold the development of our human Periportal area organoids (hPPAo) (**Materials and Methods, Figure 1c)**.

**Figure 1:**
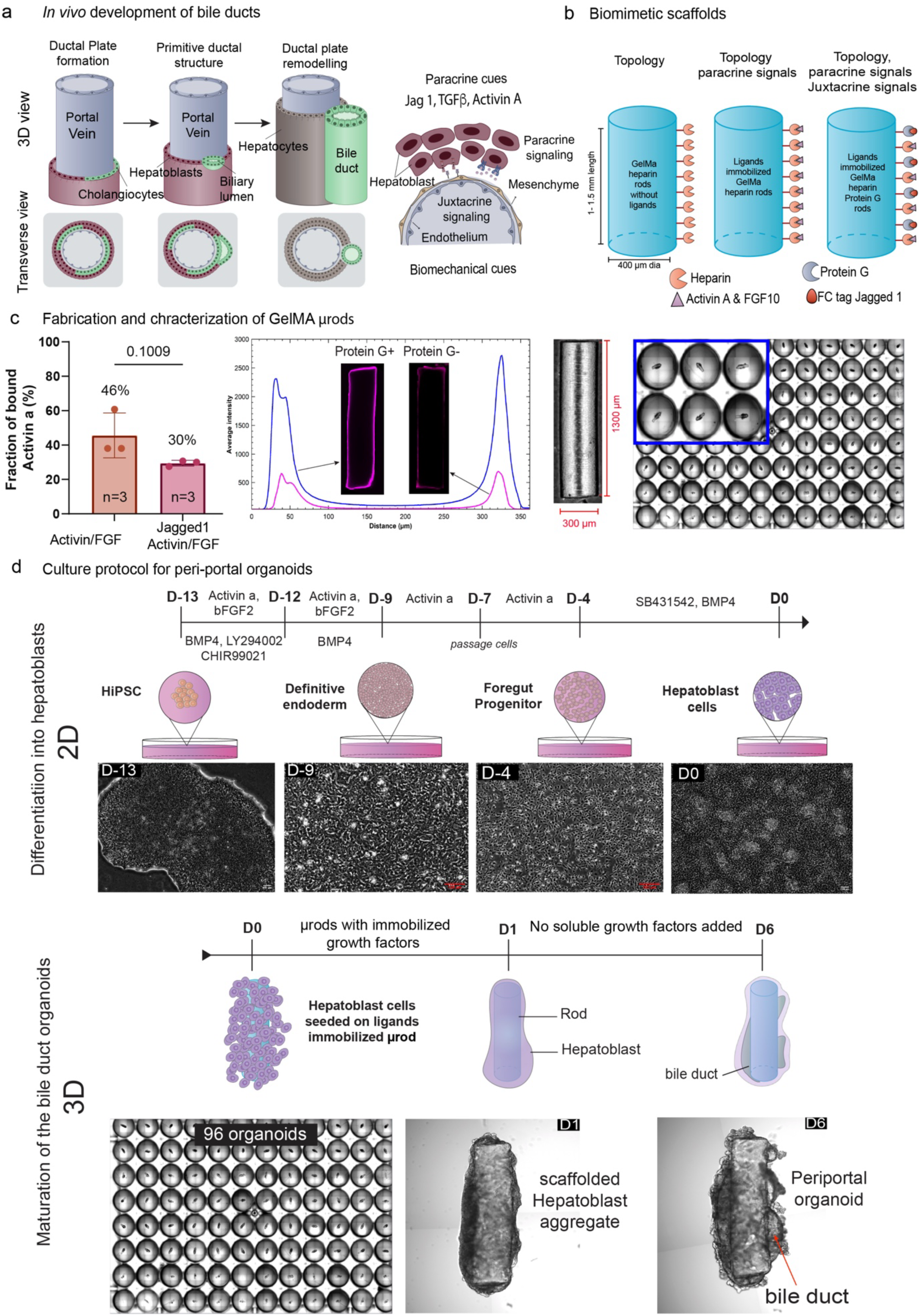
**a.** Model of bile duct development in mouse and presumably in human **b.** Schematic representation of the functionalization of the rod bio scaffold **c.** Characterization of the rod biofunctionalization in terms of grafting efficiency of Activin A and its localization on the rod. Phase contrast image of a single GelMa rod and overview of a 96 well plate with a single bio scaffold per well. **d.** Culture protocol from hIPSCs differentiation (2D) to the maturation of the hPPAo (3D).

To derive the hPPAo from hIPSCs, we first generated embryonic hepatoblasts from hIPSCs following a previously established protocol by Sampoziotis *et al* 2017, as summarized in [12]. **Figure 1d** (**Materials and Methods**). After 13 days of stepwise differentiation the vast majority of cells in the 2D culture expressed liver hepatoblasts markers (AFP and HNF4A), with a small fraction additionally expressing cholangiocyte markers including Cytokeratin 7 (CK7) and SOX9 (**Supplementary figure 1c**). We further tested the bi-potency of the hepatoblasts by differentiating them into hepatocytes and cholangiocytes on adherent culture following [25] (**Supplementary figure 1d**). The cell differentiation protocols are tuned to produce a majority of hepatoblasts, and a minority of other liver cells precursors from mesodermal lineage. This balance proves paramount to the further development of the periportal area organoids.

To initiate hPPAo formation, we harvested as single cell the 2D culture and seeded them on the biofunctionalized scaffolds in 96-well low attachment plates at a density of 50,000 cells per rod (**Figure 1d**). The seeding day sets the reference time point of the protocol Day 0 (D0). This optimized seeding density ensures a complete coverage on of the rod surface by several cell layers, without extensive cell death and the presence of a single organoid per well (**Material and Methods**). It was key to the success of the further development of the organoid.

From D0 onwards, the hPPAo grew in a maintenance medium (RPMI medium) deprived of any soluble growth factors or ECM components. We hence guided the hPPAo maturation and organisation exclusively by the biophysical and biochemical cues provided by the scaffolds as well as by the factors (growth factor and extra cellular matrix) self-secreted by the cells. In the rest of the paper, we study three combinations of biofunctionalization: the rods without any Growth factors, or biofunctionalized with Activin A and FGF10 or with Activin A and FGF10 and Jag 1.

As differentiation progressed, cells spread uniformly covering the rod surface by D1. After 6 days, the hPPAo developed mm sized elongated lumens (**Figure 1d** (D6)). The tubular structures exhibited a pulsatile behaviour (**Supplementary Video 1**), indicative of ion secretion activity into the luminal cavity.

### *In vivo* like maturation of elongated bile ducts in hPPAo

We immuno-stained hPPAo grown on rods biofunctionalized with Activin A and FGF10 for early cholangiocytes markers (Sox9, E-cadherin) and the hepatoblasts marker HNF4a. Confocal imaging (10X objective) on day 1 (D1) revealed a continuous, multi-layered coating on the rods reaching a thickness of 4 to 6 cell layers (**Figure 2a).** The innermost layer in contact with the rod segregated into two distinct populations of complementary discontinuous monolayers comprising either cholangiocytes-like cells (Sox9+, E-cadherin+) or the hepatoblasts (HNF4a+) (**Figure 2a**, **Supplementary** Figure 2a). Surrounding this interface, we observed a large fraction of HNF4a cells extending radially outwards up to 20 cell width from the rod, resembling the embryonic parenchyma. This spatial organisation is much akin to the development of the ductal plate around the portal vein (Week 8 of dev) surrounded by the embryonic parenchyma [1, 5].

**Figure 2:**
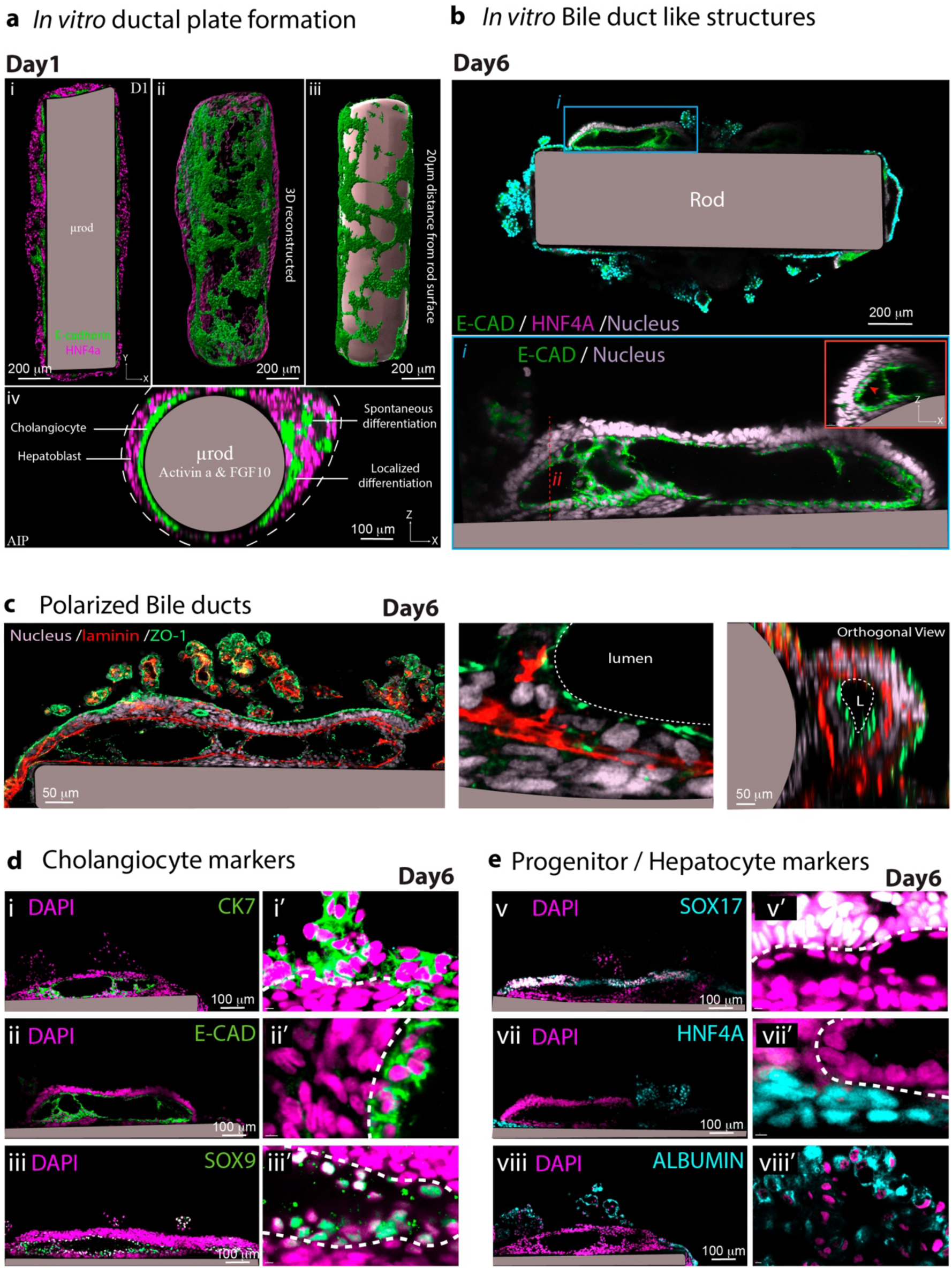
**a.i** Single-plane confocal image of the cellular distribution at Day 1 along the bioscaffold (μrod). Cholangiocytes (marked by E-cadherin) line the rod and are sheathed by hepatoblasts (marked by HNF4α) **ii** 3D volume rendering of the cellular distribution **iii** Selective 3D representation of the cell distribution within 20 μm of the rod surface **iv** Transversal view of the cellular distribution **b.** Day 6 Single-plane image of the cellular distribution at Day 6 with a close-up of the bile duct luminal cavity **c.** Day 6 single-plane image showing laminin deposition forming a basal layer for the bile duct **d.** Day 6 immunostaining for various cholangiocyte markers **e.** Immunostaining for various hepatoblast and hepatocyte markers.

Note, however that sparse cholangiocytes differentiation also occurred at distant locations away from the rod, particularly in regions with thicker parenchymal density. This “un-scaffolded” differentiation likely resulted from the presence of cholangiocytes precursors in the initial hepatoblast population at D0 following the 2D culture (see **Discussion**).

Together, our findings suggest that localized juxtacrine differentiating signals (Gelma rods + Activin A + FGF10) provided by the rods are sufficient to faithfully phenocopy the early stages of ductal plate formation.

Unlike typical *in vitro* cystic bile ducts comprising a single layer of cholangiocytes [12–15], the BEC self-organised into tubes between the rods and the surrounding parenchyma at Day 6. Immunostaining confirmed that these were *bona fide* bile duct with cholangiocytes forming the luminal lining expressed E-cadherin, CK7 and SOX9 while downregulating the progenitor markers HNF4a, Albumin, and SOX17 (**Figure 2b-c**). The lumens displayed varying morphologies including both elongated lumens and compartmentalized tubular segments (**Figure 2b**). Cells in the outer tubular layer were cuboidal with lateral E-cadherin expression (**Figure 2b**) while those in the transverse layers were more elongated (**Figure 2b** ).We further observed a clear apico-basal polarity (apical localization of the tight junction zonula occludens-1 (ZO-1) and ciliary localizing proteins ADP ribosylation factor like GTPase 13B (ARL13B)) of the ductal epithelial cells, with a continuous laminin rich basal layer delineating them from the surrounding parenchyma (**Figure 2e**, **Supplementary figure 2b**). Other transport proteins such as sodium potassium ATPase and secretin receptors (**Supplementary** Figure 2b) also polarized apically, confirming the secretory activity of the bile ducts.

### Day6 cellular diversity recapitulates that of the liver peri-portal area

To complement immunolabeling-based cell identification, we compared single cell RNA sequencing (10000 cells, approximately 5000 genes) (**Material and Methods**) at D0 (2D culture pre-seeding) and D6 (hPPAo grown of rods biofunctionalized with Activin A and FGF10).

We mapped the cell types to embryonic liver cell types using gene lists (**Material and Methods**) from based on the reported data [26–30] at a resolution of 12 clusters (**Figure 3a-b**). Single cell transcriptomics revealed four transcriptionally distinct hepatoblast clusters : eHB enriched for early markers (EFNB2, TTC6, SERPINA1), mHB expressing migratory markers (GATA4, CDH2, FN1, and GPC3), pHB other undergoing proliferation, and fHB characterized by strong expression of cell-cell adhesion proteins (CDH1) as well as foetal markers such as ONECUT2, HHEX, PROX1, and NPW (**Figure 3a-b**). From these hepatoblast, emerged the liver epithelial cells the cholangiocytes and the hepatocytes (**Supplementary figure 3a-b**). Notably, the cholangiocytes cluster BEC expressed several matrix remodelling proteins (VIM, COL1A1, COL2A1, COL3A1, LAMB1, CTHRC1, and LUM), suggestive of remodelling stages (**Supplementary figure 3a**). Supporting this, gene ontology studies revealed significant enrichment for biological process associated with epithelial tube morphogenesis (**Supplementary figure 3d**). In parallel, the hepatocytes HP expressed a spectrum of metabolic and developmental genes with the most mature ones expressing high levels of genes associated with lipid metabolism (APOA, APOB, APOE, APOC), blood coagulation (F10, FGA, SERPINA6, SERPIND1), detoxification (GSTA1, GSTA2), glucose and fatty oxidation (PDK4, CPT1A, and SLC27A5), plasma protein production (TTR, AHSG), fatty acid metabolism (FABP1) and oxidation reduction process (CYP2D10, SORD, HSD17B2) (**Figure 3a**, **Supplementary figure 3a**). The most fetal hepatocytes fHP expressed similar metabolic functions but still retained their fetal features such as PROX1, HHEX and DLK1. Despite the maturation, immunostaining did not reveal the presence of any bile canaliculi, suggesting that the hepatocytes have not reached their polarized maturation stage (**Supplementary figure 3a**). The degree of immaturity of Cholangiocytes was confirmed by the lack of expression of ABCB1(MDR1) and ABCC2(MRP2), and, for hepatocytes, the expression of cytochrome p450 genes (**Supplementary figure 3a-c**). Besides the endoderm derivatives, the hepatic stellate cells also resolved into two subclusters among which one is in activated state aHSC expressing (COL1A1, COL3A1, COL6A3, ITGA8, MMP2, ZEB2, EPHA3, MFAP4, and TNNI1) and pHSC in proliferative state (UBE2C, CDC20, CENPF, and SOX17). Endothelial cells expressing PECAM, KDR, GNG11, CDH5 and NOTCH1were also present indicating the presence periportal mesodermal cell types in the organoids.

**Figure 3:**
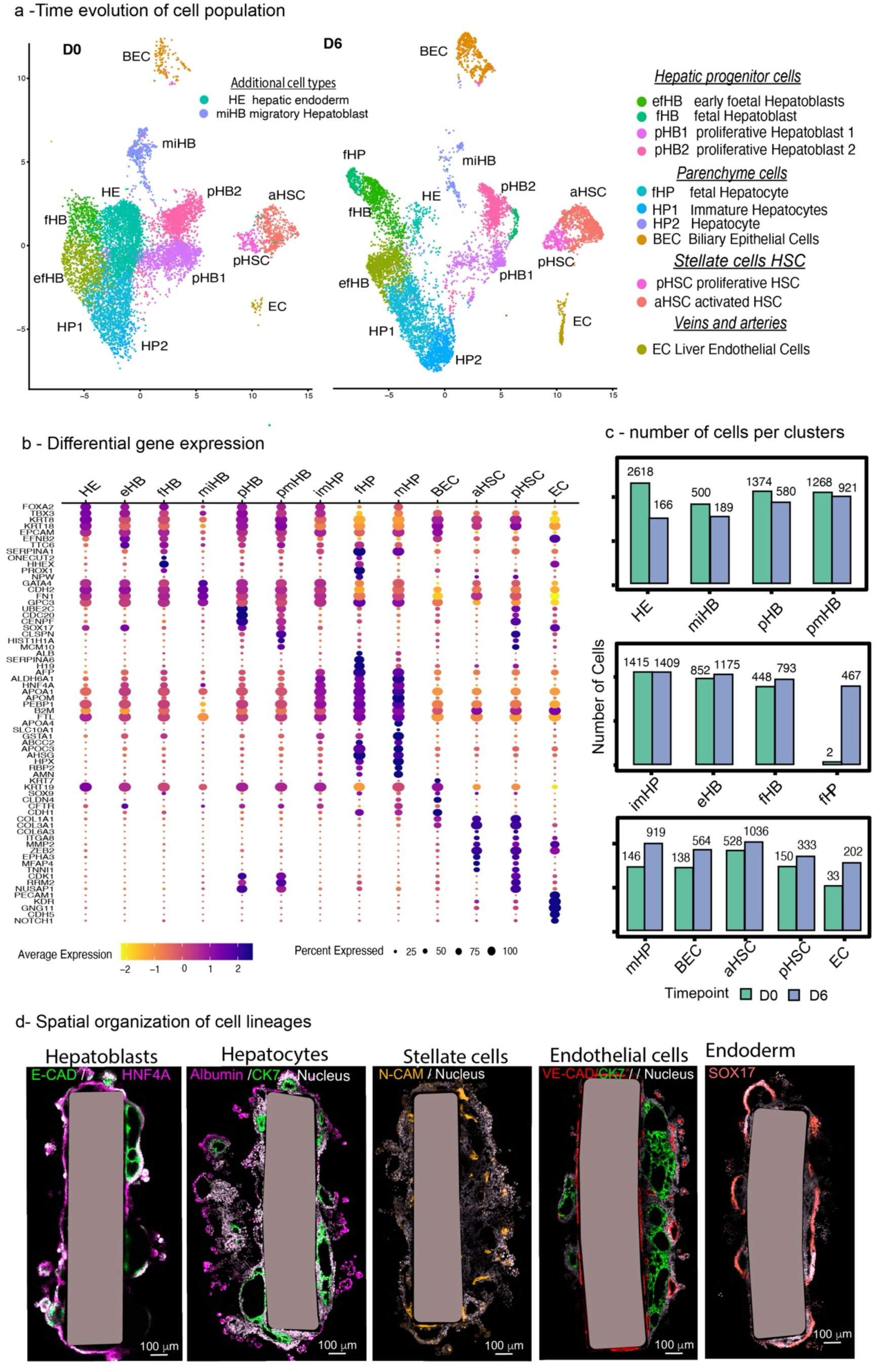
**a.** 3’ 10X single-cell RNA-seq of hPPAo at D0 and Day 6 **b.** Log2 fold-change expression levels of the principal genes overexpressed in each cell type. **c**. Number of cells per cell type at Day 0 and Day 6. **d.** D6 Single-plane confocal image showing the principal marker for each cell type; note the localization of endothelial cells as the first monolayer around the rods and as lumens adjacent to the bile ducts.

The transcriptional profile of cell types at D0 just after the phase of 2D culture (**Figure 3c**) was comparable to that of D6 although with different the relative proportion between cell types and different maturation stages (**Figure 3f, Supplementary figure 4d**). It reflected that the cellular diversity largely originated from the tuned conditions of the 2D culture steps. The D0 hepatic endoderm disappeared at day 6 (fHP, fHB) with the expansion of Hepatocytes (HP) and Cholangiocytes (BEC) as demonstrated by pseudo time analysis (**Supplementary** Figure 3b). Stellate cells matured and Endothelial cells extended. These shifts in populations occurred in the absence of any exogenous soluble growth factors and that the maturation results exclusively from self-organized secretion of differentiation factors and extracellular matrix.

We selected the most discriminative genes of each cluster and used them as ident markers to further the probe cellular spatial organization by immunofluorescence confocal imaging. Elongated bile duct and the hepatoblasts l predominantly been localized in the vicinity of the rods (**Figure 3 d**). Whereas albumin-positive hepatocytes localized preferentially at the periphery of the organoids (**Figure 3d**). The hepatic progenitor population, SOX17^+^ endoderm, consistently lined the bile duct lumens, with some proliferating sub-populations identified by Ki-67 (**Figure 3d**). Hepatic Stellate Cells (NCAM^+^) were distributed across the parenchymal regions occasionally capping bile duct lumens (**Figure 3d**). The endothelial cells (VE-cad^+^) distributed into two distinct regions: either as a layer surrounding the rods or as an epithelium lining oblate lumens. The first population can be found intercalated between the rods and the elongated bile duct, positioned on the other side of the laminin rich basal membrane, hence displacing the bile ducts outwards from the rod surface. The second population (VEGF^+^ **Supplementary figure 4a-b**) formed epithelial luminal structures in the parenchymal regions, often adjacent to the bile ducts. Together these spatial arrangements shared strong morphological resemblance with the *in vivo* portal triad configuration. (**Figure 3d**). Both populations displayed transcriptional markers for veins and arteries alike. However, they were indistinguishable from their transcriptional profile, suggesting a common progenitor identity. We did not observe the expression of STAB2 and CLEC4G which are markers of sinusoid endothelium. The induction of endothelial population in our system could originate from paracrine VEGF signalling from cholangiocytes, reminiscent of the *in vivo* developmental process. While cholangiocytes actively expressed angiogenic factors (VEGFA and ANGPT1), other cell types equally contributed to their bulk concentration (**Figure 3b, Supplementary figure 4c**). Interestingly, the corresponding receptors (KDR and FLT1) were only restricted to the endothelial clusters, suggesting a spatial organisation based on paracrine signalling and not large scale growth factor gradient (**Figure 3b, Supplementary figure 4d)**. Unexpectedly, the endothelial cell expressed the endoderm marker SOX17 instead of the mesoderm markers. Similar expression profile was already reported for hIPSCs derived liver organoids and *in vivo [31]*.

It results that the hPPAo emerge from a series of finely tune precursors in the 2D culture that mature spatially around the scaffold to give rise to a spatial cell organisation that dynamically follows the steps of periportal area maturation: development of a ductal plate and biliary tubule expansion, accompanied by the maturation of the vascular system and the parenchyma.

### Juxtacrine signalling from the scaffold governs bile duct phenotypes

We next investigated how Notch and TGF-β signalling affects the morphologies of bile ducts in hPPAo. Both pathways have been extensively studied for their roles in the bile duct developmental process [5, 32–37]. While TGF-β is known to regulate biliary differentiation and Notch acts downstream to drive bile duct morphogenesis, conflicting results exist as perturbation experiments using small molecules produce off-target effects in multicellular systems [3, 37–39]. Moreover, how these signalling pathways can be spatially controlled in *in vitro* systems to guide tissue organisation is also under active investigation.

We thus compared the developmental outcomes of the hPPAo under geometrical and ECM scaffolding only (GelMA rods without growth factors) or in the presence of paracrine differentiating growth factor signals (AF rods : GelMA rods biofunctionalized of Activin A and FGF10) or adding more morphogenetic signalling (JAF rods: GelMA rods functionalized with Jag 1, Activin A and FGF10).

At D1, in the absence of biofunctionalization (no growth factors on the rod), cholangiocyte differentiation occurred homogeneously in the parenchymal region (hepatoblasts) surrounding the rods (**Supplementary figure 5a**). Considering exclusively the first cell layer at the rod interface, reveals a sparse differentiation of hepatoblasts into cholangiocytes (**Figure 4a, Supplementary figure 5b**). As described in **figure 2a**, functionalization of Activin A and FGF10 on the rod surface enhanced the local differentiation of cholangiocytes into a discontinuous monolayer (**Figure 4a)**. Strikingly, the immobilization of Jag 1 (Notch signalling) to Activin A and FGF10 further enhanced the localization of the differentiation at the interface resulting in the formation of a continuous sheath around the rod (**Figure 4a**, **Supplementary figure 5c**). Addition of Jag 1 helped the localisation and the structuration of the morphogenesis of the organoids. **Figure 4a-b** show the quantification of the surface differentiation using a radial projection of the image of the first cell layer (**Supplementary figure 5a**) at the rod surface. The results indicate an increase in the differentiation levels from 34% in control samples to 46% in Activin A and FGF10 reaching up to 61% in Notch activated samples (**Figure 4b**). Moreover, the introduction of notch pathway reduced the number of discrete cholangiocyte clusters and increased their average area (**Figure 4b**).

**Figure 4:**
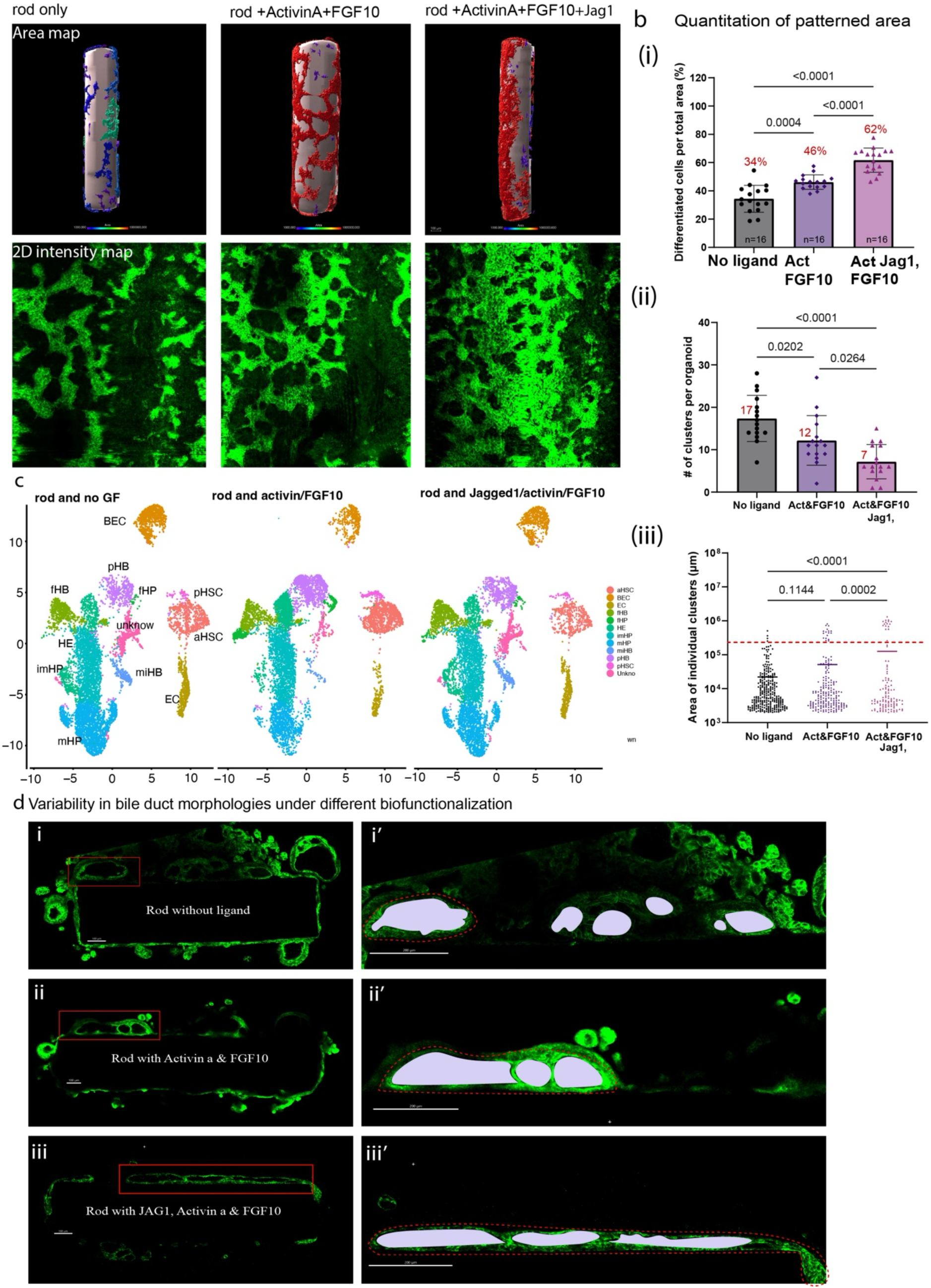
**a.** Dependence of the degree and spatial organization of cholangiocytes at Day 1 on the juxtacrine signaling environment (no growth factors on the rods, Activin A + FGF10, or Activin A + FGF10 + Jag 1). 3D volume rendering of cholangiocytes in the first cellular layer on the rod surface. Cylindrical unwrapping of raw intensity confocal images **b.** Quantification of morphological signatures derived from the cylindrical projections: **i** fraction of cholangiocytes in the first monolayer **ii** number of disconnected clusters **iii** area of individual clusters **c.** UMAPs from 3’ 10X single-cell RNA-seq comparing cell populations under each juxtacrine signaling condition, showing no significant differences **d.** Variability of bile duct morphologies under different juxtacrine signaling conditions at Day 6; rods without growth factors generate discontinuous spherical lumens, Activin A + FGF10 induces longer, compartmentalized tubular ducts, and Activin A + FGF10 + Jag1 generates long, connected tubular ducts spanning the entire rod length.

At D6, UMAP clustering of scRNAseq data comparing all three conditions (**Figure 4c**) did not reveal any measurable difference in term of cellular composition. This result further suggests that the presence of grafted growth factors on the rods does not influence the cellular diversity that is likely to originate from the self-induced differentiation of the original cell population at D0. In particular, the cholangiocytes across all conditions expressed key markers such as Cytokeratin 7, SOX9 and E-cadherin, with a concurrent reduction in hepatoblast and hepatocyte marker albumin (**Supplementary figure 6**). It provides guiding cues to enhance the maturation and structuration programs of the cellular types. Notably, the induction of Notch led to the expression of MDR1, an ATP-dependent efflux pump crucial for cholangiocyte protection, suggesting a role for notch signaling in cholangiocyte maturation (**Supplementary figure 6a**). However, the bile ducts were unable to transport the bile acid analogue, Cholyl lysyl fluorescein (CLF), suggesting they have not achieved full functional capability (**Supplementary figure 6a-b**). This result is expected since the developmental stage we mimic is around Week 14-15 whereas the bile acid transport of cholangiocytes only starts around Week 19-20 in humans.

Despite similar cellular composition across different conditions, hPPAos presented salient differences in bile duct morphology. In control samples, cholangiocytes self-organized to form oblate lumens with minimal elongation (**Figure 4d**). With Activin A and FGF10 bile ducts appeared as elongated interconnected lumen that maintained intact epithelial walls between adjacent structures (**Figure 4d**). Notably, the presence of Jag 1 enhanced this remodelling process, characterized by the coalescence of lumens into long continuous ducts along the rod axis (**Figure 4d**). This progressive tubular formation, achieved with minimal fusion events is reminiscent of *de novo* bile duct morphogenesis. Our results confirm *in vitro* the direct influence of Notch signalling on the ductal plate remodelling.

### Quantification of bile duct tubulogenesis by Optical coherence tomography

The assessment of the bile duct morphologies using confocal microscopy (Material and Methods) required prolonged imaging (more than 10 hours per organoid at 10X objective) yielding a very low throughput and limited imaging depth (typically up to 200μm in the hPPAo). Although distinct lumen morphologies were distinguished visually, the method lacked standardized pipelines for unbiased detection and quantitation. To overcome these limitations, we used Optical Coherence Tomography (OCT), a label-free imaging technology that leverages on refractive index changes between the lumens and the surrounding tissues to generate 3D high contrast images (**Figure 5a-b**). OCT offers sufficient depth penetration (up to to 2mm) in thick tissues with high spatial (4.8 μm) and temporal resolution (10 s for *in toto* imaging of the organoid). It hence enabled rapid 3D volumetric imaging of lumens. (**Supplementary figure 7**).

**Figure 5:**
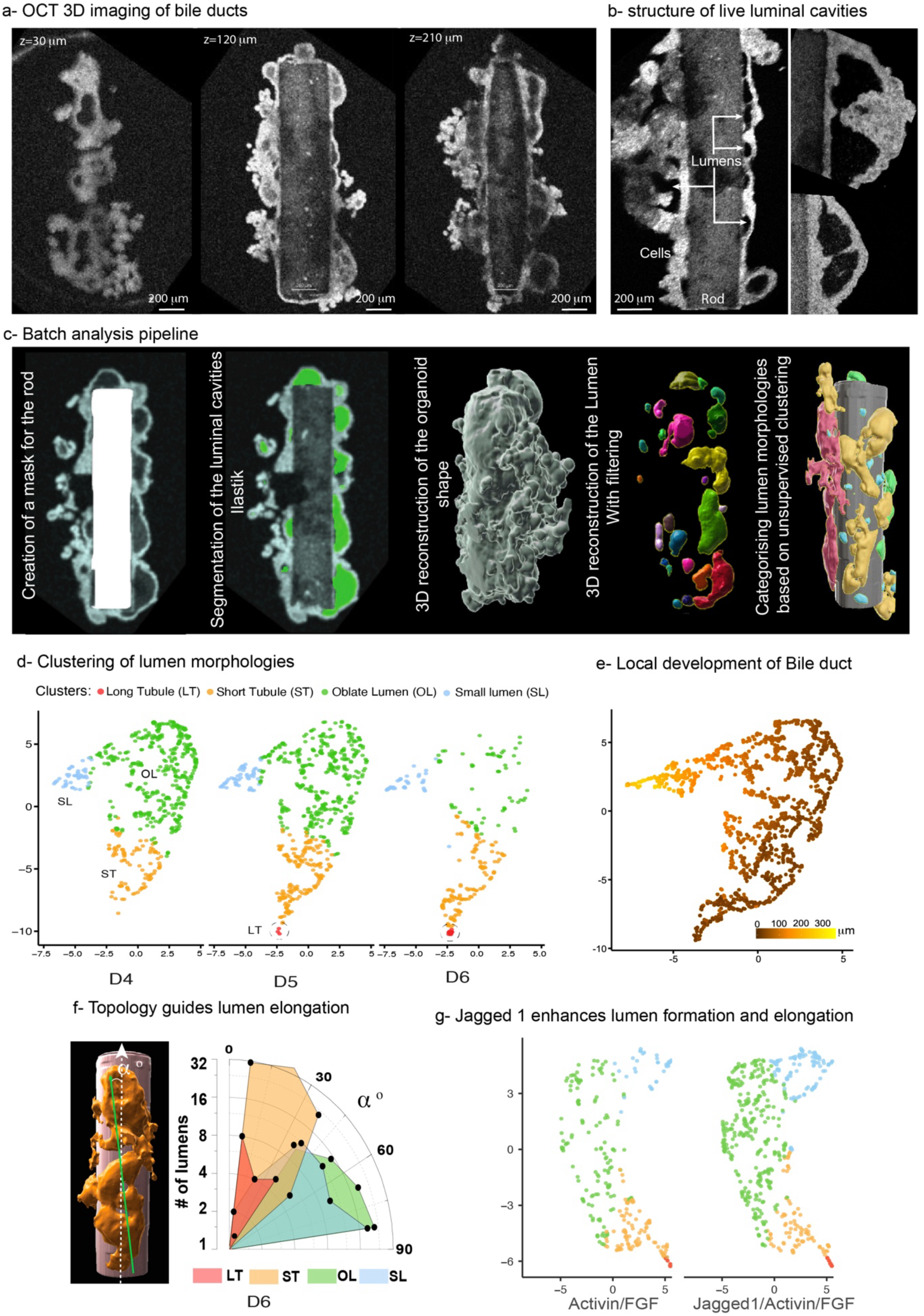
**a.** Optical sections of a 3D hPPAo using Optical Coherence Tomography (OCT); the cellular, rod, and luminal regions are clearly distinguishable **b.** High-resolution OCT close-up of cellular septa within elongated tubular bile ducts (indicated by yellow arrows), indicative of connections between different compartments **c.** Schematic of the batch analysis process: establishment of a mask for the rod interior, training and detection of luminal cavities using Ilastik, 3D reconstruction of the hPPAo volume, 3D segmentation of individual lumens, followed by unsupervised clustering of lumen morphologies **d.** Clustering (UMAPs) of seven morphological features of lumens comparing Day 4, Day 5, and Day 6; the long bile ducts (around one per organoid ), appear as a small distinct cluster (red) **e.** Heat map of the distance of each lumen cluster relative to the rod surface; longer lumens tend to be located closer to the rod **f.** Radial histogram of the distribution of angles between the long axis of the ducts and the rod axis for each cluster type; although the angle contributes to clustering, elongated lumens consistently align with the rod’s main axis **g.** Clustering (UMAPs) of seven morphological features of lumens comparing Activin A + FGF10 and Activin A + FGF10 + Jag 1 conditions; Activin A + FGF10 + Jag 1 leads to an overall increase in the number of lumens, with a higher proportion of long tubular structures

We performed extensive imaging of 96-well plates on a custom-made OCT imager (**Materials and Methods**). To extract and to analyse the lumen morphological features, we developed a semi-automated image processing pipeline as detailed in the supplementary information (**Supplementary** Figure 8). Briefly (**Figure 5c**), the raw OCT images were subjected to simple pre-processing schemes including denoising and contrast enhancement. A custom developed algorithm (**Material and Methods**) was then employed to extract the mask of the rod volume by coupling water-shading and a-priori knowledge of the rod shape. Using sparse 2D planes of the organoid images, we then trained Ilastik (running on an ImageJ environment) to segment the lumen cavities from the cellular areas. Batch processing for the entire z-stacks of individual organoids was then possible. However, due to the experimental variability of the imaging noise, the training was performed for each imaging batches. Following segmentation, a 3D volume representation of each hPPAo was obtained from the segmented image stacks in IMARIS (Andor). We further extracted morphological features of each segmented luminal cavities – Sphericity, Volume, Shortest distance to the rod, Elipsoid long axis length, orientation angle - using IMARIS analysis modules (**Figure 5c**). The smoothing parameters to merge adjacent lumens were adjusted by comparison of the raw images to the segmented ones and were kept consistent across all hPPAo batches.

We then performed k means clustering on 4 principal morphometric parameters (sphericity, volume, shortest distance to the rod and ellipsoid long axis length) of the lumens to classify them in an unbiased manner (**Supplementary figure 9**). This was first performed with pooled datasets from three developmental time points (D4, D5, D6) for rods biofunctionalized with Activin A and FGF10. The unsupervised clustering revealed 4 different groups: SL for Small Lumens. OL for Oblate lumens, ST for Short Tubes and LT for long Tubes named to reflect the dominant geometry within each group. Of these the LT, ST and OL clusters were predominantly localized proximal to the rod surface with their average distance ranging ∼25µm **(Supplementary figure 9biii)**. The LT cluster displayed features of bile duct morphology with tubes exhibiting a sphericity of 0.44 and volume of 94,00000 µm³ with length extending almost 7 to 8 times longer than lumen clusters **(Supplementary figure 9b).** These tubes were typically found at a frequency of 1 per organoid. Located significantly distant from the LT clusters are the ST clusters that exhibited smaller, non-spherical lumens with an average volume of ∼25,0000 µm³ and a sphericity of 0.8 **(Supplementary figure 9b)**. These were predominantly populated by either spontaneously differentiated cholangiocytes or endothelial cells, as evidenced by CK7 and VE-cadherin staining, respectively **Supplementary figure 4a).** It was not possible for us to morphologically discriminate the small lumens from endothelial cavities using OCT images.

Temporal analysis further revealed the progressive development of long tubes from D5 and the gradual loss of oblate lumens over time, correlating with a decrease in the overall sphericity and an increase in volume as tubes formed within organoids (**Supplementary figure 9b**). This corroborates with the idea of a gradual fusion of individual cystic bile ducts into a single long tube. This transition was further characterized by the high resolution (20μm) OCT live images (**Figure 5a**) where we detected the presence of small gaps between adjacent luminal cavities indicate of active fusion events. Taken together, OCT imaging in combination with unsupervised clustering enables the identification and temporal tracking of lumen morphologies including long bile ducts in hPPAo.

We then computed the angle between the rod axis and the principal axis of each D6 lumen (**Figure 5e**) to assess the level of alignment of tubular structures with the rod scaffold. Lumen in LT and ST clusters showed a clear orientation along the rod axis whereas OL and SL were more isotopically distributed with some biased orthogonal to the main axis. Notably, orientation angle was not used as a clustering criteria validating the robustness of the observed pattern. These findings indicate that the geometrical cues provided by the rod orient the development of the bile duct directionally along its axis rather than spiralling around the rod. To validate this hypothesis, we performed experiments wherein we provided a different topology in the form of beads (µbeads), while maintaining the same paracrine cues - Activin a/FGF10 **(Supplementary figure 10a)**. Despite the changes in the topology, Cholangiocytes differentiated locally near the bead surface, as evidenced by SOX9 and E-cadherin expression **(Supplementary figure 10b)**. Similar to the rod system, the differentiated cells formed localized lumens consisting of a single layer of cholangiocytes with the proper establishment of cell polarity **(Supplementary figure 10b&c)**. However, even after prolonged culture, these organoids did not progress into tubes, strongly supporting the role of portal vein topology in guiding the bile duct tube formation (**Supplementary figure 10b**).

We finally performed k means clustering to compare Activin A/FGF10 and Jag 1/Activin A/FGF10 conditions (**Figure 5g, Supplementary figure 11**). It confirmed the existence of 4 clusters that matched the characteristics of those found previously. They key difference is the number of lumens detected per cluster with an increase in the number of long tubes in Jag1/Activin A/FGF10 activated rods **(Supplementary figure 11).**

## Discussion

Cholangiocyte organoids have been generated in many instances, with a strong emphasis on cholangiocyte maturation and less attention given to tubulogenesis. In most cases, spherical cholangiocytic cysts are produced by embedding primary or hIPSC-derived cholangiocytes in a 3D extracellular matrix (ECM). Optimizing the matrix properties can enable the formation of branched ducts from adult (but not fetal) primary cholangiocytes [23]. Most existing organoid models focus primarily on bile duct maturation. To achieve greater cellular complexity— including blood vessels, stellate cells, smooth muscle cells, and hepatocytes—the common strategy has been to derive these cell types separately and combine them into a single organoid that is further matured through transplantation. Recent work [24] demonstrated that hIPSC-derived cholangiocytes can be cultured on hIPSC-derived vascular cells preorganized into tubular structures. This optimized protocol showed that the presence of an artificial blood vessel promotes the development of elongated bile ducts, which can subsequently be transplanted into mice for long-term maturation. Recently mouse periportal assembloids have also been derived by the re-association of mouse purified hepatocytes, cholangiocytes and portal mesenchyme in carefully selected proportions [40]. Such an approach is not readily adaptable to human to produce large number of organoids. Our work proposes an orthogonal method. We first established a protocol to grow hIPSCs with a diversity of liver-associated cell types. We then designed an ECM-biofunctionalized scaffold to coordinate the organization of these cell types into complex organoids that not only recapitulate bile duct elongation but also mimic the periportal area. This occurs through the self-organization of diverse liver cell populations and the reproduction of key physiological steps of bile duct development, from ductal plate formation to duct fusion. Although lacking specific arterial or venous markers, the endothelial structures organized in a stereotypical manner, eventually occupying the space between the rod and the bile duct and forming luminal structures around it. In its current form, the organotypic assembly also displays a heterogeneous parenchyma containing hepatic precursors, immature hepatocytes, and stellate cells. Our hPPAo system differs from existing models in that it is highly scalable (no bioprinting required, compatible with 96-well plates) and optimized for quantitative imaging. The scaffolds here very simply replace the biophysical guidance of the portal vein as well as the chemical juxtacrine signalling to elicit local differentiation. It offers a reductionist, highly tunable technique to study periportal tubulogenesis. It offers a first example but can be adapted to other organoid types or complexify to create gradient of growth factors leading to zonation. However, as a seminal step, this paper focuses on the first seven days of differentiation, corresponding to early morphogenetic events (equivalent to weeks 12–16 in human development). It offers a first example and can be adapted to other organoid types or complexify to create gradient of growth factors leading to zonation. However, as a seminal step, this paper focuses on the first seven days of differentiation, corresponding to early morphogenetic events (equivalent to weeks 12–16 in human development). Further maturation and transplantation are beyond the scope of this study.

This bioscaffold approach is scalable and compatible with parallel culture formats such as 96-well plates. We leveraged this by developing a semi-automated screening method using Optical Coherence Tomography (OCT) to classify luminal cavities. This work serves as a proof of concept demonstrating how scaffolded hPPAo culture, OCT imaging, and quantitative analysis can be integrated into a pipeline. Further work is needed to fully automate image acquisition, as the current analysis still depends on experimentalist input. One area for improvement is the development of an objective criterion to determine whether adjacent lumens are connected, which remains a limitation of our current analysis and requires experimental validation. Since high-resolution OCT imaging provides clear visualization of connecting gaps, we are developing AI-based methods to recognize such features in 3D, enabling reliable discrimination between adjacent and truly connected bile ducts. Enhancing image contrast consistency across batches and training Ilastik on larger datasets will also reduce the need for frequent retraining of the lumen segmentation algorithm.

## Conclusion

The portal vein biomimetic scaffold approach to hepatic organoids enables faithful recapitulation of the early steps of periportal area formation, including bile duct elongation, vascularization, and mesenchymal maturation. This strategy is based on generating a mixed population of liver cell precursors in 2D culture and using a bioscaffold to guide their self-organization and maturation. Through this system, we identified the role of geometrical cues in aligning duct elongation along the scaffold axis, as well as the role of growth factors (Jag 1 and TGF-β) in promoting the fusion of immature lumina into millimeter-scale bile ducts. The method is both scalable and optimized for quantitative imaging using optical coherence tomography. Together, these features make the approach a powerful platform for scaffold-based organoid development.

---

**Supplementary Figure 1: a.** Schematics of µrod fabrication using a scalable extrusion method. **b.** Biophysical characterization: rod dimensions (n=33 samples) **(i)** Elastic modulus of scaffolds using AFM **(ii)** Concentration of Activin A immobilized per rod measured using ELISA **(iii)**. **c.** Immunofluorescence staining of D0 hepatoblasts with its characteristic marker (HNF4A & AFP) and cholangiocyte markers (SOX9 & CK7). Scale bar = 200 µm. **d.i** 2D differentiated hepatocytes displaying cuboidal morphology and expressing ASGR1(blue) and HNF4A (red) markers. **d.ii** Cholangiocyte progenitors embedded in 3D matrigel showing lumen formation and stained for CK7 and AFP.

**Supplementary Figure 2: a.** Characterization of cholangiocyte differentiation in early-stage organoids. Immunofluorescence staining of D1 organoids showing SOX9^+^ **(i)** and E-cadherin^+^ **(ii)** cholangiocytes surrounded by HNF4A^+^ hepatoblasts **(iii)** with composite image of the two population **(iv)**. Scale bar = 5µm. **b.** Traverse images of organoids showing key cholangiocyte polarity proteins: apical ARL13B **(iii)**, basolateral Na^+^K^+^ ATPase **(iv)**, diffused SCTR **(v)** and basal laminin **(i)** scale bar = 100µm. Insets **(i’,ii’,iii’,iv’)** show magnified images with red arrowheads indicating localization. Dotted lines indicate the location of bile ducts. in magnified images represent the localization position of proteins.

**Supplementary Figure 3: a.** Dotplot showing metabolic gene expression in hepatocyte cluster. **b.** Pesudotime trajectories demonstrating the emergence of biliary epithelial cells, mature hepatocytes and fetal hepatocytes. **c.** UMAP overlay showing expression of ABCC2 (MRP2). **d.** Volcano plot highlighting the differential gene expression between D6 biliary epithelial cells and mature hepatocytes.

**Supplementary Figure 4: a.** Transverse images of D6 organoids showing spatial organization of endothelial (VE-cadherin, red) and bile duct lumens (CK7, green). **b.** Dotplot showing the expression of various vasculature associated genes in D6 organoids. **c.** Violin plots showing the expression of ligands VEGFA, ANGPT1 in the cholangiocytes and other clusters and its corresponding receptors KDR, FLT1 in the acceptor cell, endothelial cluster. **d.** Expression of endoderm marker SOX17 in endothelial clusters. **e.** Temporal UMAPS (D0, D1, D6) illustrating the shift in cellular composition and number over time.

**Supplementary Figure 5: a.** Quantitative assessment of patterning area on rod surface. Workflow: LSM raw data → rod alignment → 2D unwrapping → generation of intensity maps → quantification of differentiated cell area. **b.** Confocal and 3D renderings of control rods showing heterogeneous cholangiocytes distributed in hepatoblast organoids. **c.** Jag 1- functionalized rods showing improved cholangiocye monolayer formation around rod surface. Cross-sectional and intensity projection images indicate the differences

**Supplementary Figure 6: a.** Phenotypic analysis of control rods, rods with Activin A and FGF10 and rods with Jag 1, Activin A and FGF10 activation using cholangiocytes markers SOX9, CK7, E-cadherin, MDR1, as well as hepatocyte and hepatoblast marker albumin. **b.** Cholangiocyte functional assay using CLF on D6 organoids across different conditions.

**Supplementary Figure 7: a.** Comparative analysis of LSM with OCT imaging at different focal depths, revealing better depth penetration, high spatial and axial resolution with OCT. **b.** OCT images of fixed **(i)** vs live D6 organoids **(ii)** revealing better tissue contrast in live samples, thus distinguishing cells, rod, lumens and background

**Supplementary Figure 8:** Automated segmentation pipeline using machine learning tool (1) High-content image acquisition of D6 bile duct organoids using OCT. (2) Preprocessing steps to automatically rotate and align rods along vertical orientation to facilitate rod mask generation on IMARIS. (3) Generation of lumen mask with supervised training in ilastik. (4) 3D volume rendering and visualization of lumen in IMARIS. (5) Extraction of morphological parameters and downstream analysis to address various biological questions.

**Supplementary Figure 9:** Quantitative morphometric analysis of bile duct lumens across developmental timepoints **a.** UMAP colour coded for sphericity (i) and long axis length (ii). **b.** Quantitative analysis of morphometric parameters among different categories. (i) Statistical comparisons of lumen categories across D4, D5, D6 organoids: number of lumens (i) lumen sphericity (ii), shortest distance to rod surface (iii), long axis length (iv) and volume (v). N = 10 organoids; Significance was calculated using an ordinary one-way ANOVA (GraphPad Prism).

**Supplementary Figure 10: a.** Schematic representation of the gelatin heparin bead coupled with Activin A and FGF10 growth factors. **b i-iii’.** Phenotypic and morphological changes in bile duct organoids, stained with SOX9 (magenta), E-cadherin (green) and Actin (yellow). **c.** Quantification of the relative SOX9^+^ cells compared to the total cell count. The colours on the graph correspond to the respective markers in b. Statistical significance was calculate for SOX9^+^ cells is indicated as follows: ns-nonsignificant, P > 0.05; *P ≤ 0.05; ** P ≤ 0.01; ***P ≤ 0.001; ****P ≤ 0.0001(one-way ANOVA).

**Supplementary Figure 11: a.** Morphometric comparison of lumen features between Activin A FGF10 rods and JAG1+Activin A FGF10 rods, including number of lumens (i), lumen sphericity (ii), shortest distance to rod surface (iii), long axis length (iv) and volume (v). n = 10. Significance tested by one-way ANOVA (GraphPad Prism).

**Supplementary video 1:** Live imaging showing expansion of lumens in hPPAo.

## Acknowledgments

VV acknowledges support from MBI core funding, CALIPSO grant NRF2019-THE002-0007, and Chaire d’Excellence AMIDEX 2ORGASRH. The hIPSCs used in this work were a kind gift from Dr. S. Tamir Rashid (Imperial College London, UK). The authors collectively thank Dr. C.T. Lim for the use of its AFM and Dr. Don Ingber for the use of the Pluto software, Dr. Andrew Holle for letting us use the live microscope. PM acknowledges support from the CENTURI Hackathon for initial trials of OCT images segmentation strategies. VV and HR also acknowledges the Pluto team for all their support and help. VV and HR acknowledges Dr. Tony Kanchanawong for his support and assistance.

## AUTHORS CONTRIBUTIONS

HR and VV designed the research. HR, MM, GS performed the biological experiments. HR, FL, SS, AM, OHT, PM performed the image analysis. HR and AA engineered the rod approach. HR and JZ performed bioinformatic analysis for all the single-cell RNA-seq data and LC helped in scRNA-seq data analysis. NMHB provided technical assistance for AFM measurements. MH, JL, FN, BK provided technical and assistance support for the OCT imaging. HR and VV wrote the manuscript.

## Material and Methods

### Fabrication and functionalization of GelHep μrods

GelHep μrods were fabricated by injecting a prewarmed precursor solution [5% (w/v) GelMA with 2.5% (w/v) heparin sulfate at a 2:1 ratio] into passivated TYGON tubing (0.5 mm inner diameter; Cole-Parmer) and UV crosslinked (50% power, 10 min). The tubings were then cut into ∼1–1.5 mm segments and subjected to slow ethanol 90% (v/v) dehydration overnight at 4 °C on shaking rocker to facilitate the release of rods. The following day, rods were collected and chemically stabilized by crosslinking in 90% ethanol with 2 mg/ml EDC and 0.48 mg/ml NHS for 6 h at room temperature before sterilizing in 70% ethanol.

For ligand coupling, the sterile rods were preactivated in 2 mM EDC and 5 mM NHS for 15 min at room temperature and incubated in Protein G solution for 1 h. Growth factors were then coupled to the rods by resuspending the sterile rods in growth factor solution (5X the concentrations used for differentiation; if Jag 1 = 20ng per rod) overnight. Control rods were incubated in PBS without ligand.

### Fabrication and functionalization of GelHep μbeads

GelHep μbeads were generated by emulsifying GelHep solution [10% (w/v) gelatin and 5% (w/v) heparin sulfate (Sigma-Aldrich) in sterile distilled water] in mineral oil containing 2% (v/v) SPAN80 (Sigma-Aldrich) at a ratio of 1:25 and stirred at low speed for 5 min. The emulsion was subsequently cooled on ice for 10 min, after which an equal volume of 30% (v/v) ethanol (Sigma-Aldrich) was added and the suspension was centrifuged at 1500 rpm for 5 min to promote phase separation. Beads were sequentially washed with ethanol solutions of increasing concentration up to 90% (v/v) and were crosslinked with EDC (2 mg/ml) and NHS (0.48 mg/ml) for 6 h at room temperature. Beads were then stored in 70% ethanol overnight for sterilization.

For ligand immobilization, μbeads (100 μl suspension) were pelleted (5000 rpm, 3 min) and resuspended and incubated overnight (4 °C) in PBS containing growth factors (5X differentiation concentration). Control beads were incubated in PBS without ligands.

### Enzyme-linked immunosorbent assay (ELISA)

Immobilization efficiency was quantified using the Quantikine ELISA kit for Activin A (R&D Systems) according to the manufacturer’s instructions. Briefly, 100 GelHep μrods were incubated in PBS containing 250 ng/ml Activin A for 24 h at room temperature and immobilized amounts were calculated from the depletion in the supernatant.

### Measurement of GelHep μrods Elastic modulus

The mechanical properties of GelHep μrods were measured using a NanoWizard IV BioAFM system (JPK Instruments, Germany) in collaboration with Dr. C.T. Lim’s group. Prior to measurement, μrods were allowed to swell in PBS for 5 min and affixed to plastic Petri dishes. Indentation measurements were performed on 18 μrods obtained from two independent fabrication batches. A polystyrene bead (45 μm diameter; Novascan Technologies, USA) was attached to a cantilever with a spring constant of 0.17 N/m, and indentations were carried out using a loading force of 5 nN at a speed of 5 μm/s.

Young’s modulus values were extracted from approach curves using JPK Data Processing Software (JPK Instruments, Germany), applying the Hertz contact model for a spherical indenter (diameter 45 μm; Poisson’s ratio 0.5).

### Protein G quantification

Protein G coupling on GelHep μrods was validated by antibody binding. Protein G–coupled μrods were incubated with rabbit secondary antibody (1:500 dilution) and washed extensively with PBS to remove unbound fraction. Fluorescence imaging was performed using an Olympus FV3000 NIR confocal microscope, and signal intensity profiles were quantified using ImageJ Intensity Profile Tool.

### hIPSC culture

Human induced pluripotent stem cells (hIPSCs; provided by Dr. Tamir Rashid, King’s College London) were maintained on Vitronectin XF–coated culture flasks (Nunc™ EasYFlask™, Thermo Fisher Scientific). Cells were cultured in TeSR-E8 medium (STEMCELL Technologies) with daily medium changes and passaged every 5 days using ReLeSR (STEMCELL Technologies). For passaging, cells were replated as small colonies.

### Differentiation protocol

#### Generation of hepatoblasts from hIPSCs

hIPSCs were differentiated into hepatoblasts following established protocols. Colonies of appropriate size were dissociated with ReLeSR (STEMCELL Technologies) and seeded onto 6-well tissue culture plates (Corning Costar) pre-coated with 0.1% gelatin and mouse embryonic fibroblast (MEF) medium [DMEM (Sigma-Aldrich) supplemented with FBS (Gibco), L-glutamine (Gibco), penicillin–streptomycin (Gibco), and 2-mercaptoethanol (Gibco)]. Cells were primed in TeSR-E8 medium (STEMCELL Technologies) for 24 h and maintained in a hypoxic incubator at 37 °C with 5% CO₂ and 5% O₂.

Hepatoblast differentiation was induced by sequential addition of growth factors and small molecules in RPMI medium (Sigma-Aldrich) supplemented with penicillin–streptomycin, MEM non-essential amino acids (MEM-NEAA, Life Technologies), and B27 supplement (Life Technologies). The following reagents were applied to target definitive endoderm, foregut, and hepatoblast stages: Activin A (100 ng/ml, R&D Systems), FGF2/bFGF (80 ng/ml, R&D Systems), BMP4 (10 ng/ml, R&D Systems), LY294002 (10 μM, R&D Systems), CHIR99021 (3 μM, R&D Systems), Activin A (50 ng/ml, R&D Systems), SB431542 (10 μM, R&D Systems), and BMP4 (50 ng/ml, R&D Systems).

#### Differentiation of hepatoblasts to cholangiocytes using ligand-immobilized μrods

Day 13 hepatoblasts were dissociated into a single-cell suspension using Accutase (STEMCELL Technologies) and were seeded (10,000 per rod) in low-attachment 96-well plates (Corning) together with one GelHep μrod per well in a total volume of 200 μl RPMI/B27 medium supplemented with Y-27632. After 24 h of aggregation, medium was refreshed daily for 6 days in the absence of soluble growth factors or Y-27632.

#### Differentiation of hepatoblasts to cholangiocytes using ligand-immobilized μbeads

For μbead induction, day 13 hepatoblasts were seeded at a 1:10 bead-to-cell ratio under the same aggregation and culture conditions as for μrods.

#### Immunostaining

3D samples were fixed in 4% paraformaldehyde for 20 min at room temperature or in a 37 °C incubator, followed by permeabilization in 1% Triton X-100 in PBS overnight at 4 °C. Samples were then incubated in blocking buffer [2% (w/v) BSA and 1% Triton X-100 in PBS] overnight at 4 °C, followed by incubation with primary antibodies diluted in antibody buffer [2% (w/v) BSA and 0.2% Triton X-100 in PBS] for 48 h at 4 °C.

Primary antibodies used were: HNF4A (rabbit, Cell Signaling Technology), AFP (mouse, Abcam), cytokeratin 7 (rabbit, Abcam), SOX17 (goat, R&D Systems), SOX9 (rabbit, Cell Signaling Technology), E-cadherin (mouse, BD Biosciences), and laminin (rabbit, Sigma-Aldrich). Samples were washed twice in washing buffer [3% (w/v) NaCl and 0.2% Triton X-100 in PBS] at room temperature for 1 h, followed by overnight incubation at 4 °C.

Secondary antibody incubation was performed for 48h at 4 °C in the presence of DAPI (0.5 μg/ml; Invitrogen) and ATTO-565 phalloidin (ATTO-TEC). Washing was carried out as described for primary antibodies. Samples were cleared in prewarmed RapiClear 1.52 (Sunjin Lab) and mounted using spacers (0.2 mm for organoids with μbeads; 0.5 mm for organoids with μrods) before sealing with a coverslip. Immunofluorescence images were acquired using an Olympus FV3000 NIR confocal microscope.

#### Live-cell imaging by bright-field microscopy

For live imaging, organoids were transferred from low-attachment round-bottom 96-well plates into low-attachment glass-bottom 96-well plates. Imaging was performed using a Zeiss Celldiscoverer 7 widefield microscope, with images acquired once per hour over a 3-day period.

#### Single-cell RNA sequencing

Single-cell suspensions were prepared from 50 organoids by dissociation with Accutase (STEMCELL Technologies) for 20 min. The aqueous layer containing cells was collected, leaving rods settled at the bottom, and passed through a 40 μm cell strainer to generate uniform suspensions. Following centrifugation, cells were resuspended in PBS containing 0.1% BSA. Libraries were prepared using the 10x Genomics platform, and sequencing data were processed with Cell Ranger to generate count matrices.

Downstream analysis was performed using Seurat v5. Count matrices were converted into Seurat objects, and quality control filters were applied to remove cells with >10,000 or <2,000 detected genes, >100,000 or <5,000 total molecules, or >10% mitochondrial reads. Doublets were identified and removed using the *scDblFinder* package. Data were normalized using the NormalizeData function in Seurat, and 2,000 highly variable genes were selected for further analysis. Datasets were scaled, and principal component analysis (PCA) was applied for linear dimensionality reduction. Clustering was performed on up to 30 PCs at a resolution of 0.5, and non-linear dimensionality reduction was performed using UMAP. Clusters were annotated manually based on expression of canonical marker genes (Table 7.1).

#### Image analysis and quantification

3D rendering was carried out using IMARIS (v10.1, Bitplane) and Fiji (ImageJ). Graphs and statistical analyses were generated using Origin (OriginLab). Single cell analysis was performed using Seurat v5 and Pluto. Lumens k means clustering was performed using R.

#### Optical coherence tomography

The cell-laden scaffold samples were imaged using a fiber-based spectral-domain OCT system (TEL220, Thorlabs Inc., USA). The focused optical beam in the OCT system illuminates the sample with nonionizing, near-infrared light with a central wavelength of 1300 nm and a spectral bandwidth of 170 nm. The OCT scan lens used (LSM02, Thorlabs Inc., USA) has a numerical aperture of 0.11. The backscattered light returning from the sample is detected on a spectrometer and the time delay between the sample and reference arms of a Michelson interferometer is used to reconstruct a depth profile (A-scan) of sample backscattering. The OCT beam was scanned laterally to acquire two-dimensional (B-scan) and three-dimensional images of the sample, with experimentally measured axial and lateral resolutions of 4.8 μm and 4.4 μm, respectively, to 1−2 mm in depth. Each sample was scanned while submerged in cell media in a well of a 96-well plate. Scans comprised 1,000 A-scans per B-scan, and 1,000 B-scans per volume over a 2×2 mm (xy) lateral field-of-view, resulting in a lateral sampling density of 2 μm per voxel. The axial sampling density, set by the specifications of the spectrometer, was 2.5 μm per voxel. The A-scan acquisition frequency was 50 kHz, resulting in an OCT volume acquisition time of approximately 20 seconds.

| Step | Description | Parameters / Thresholds |
| --- | --- | --- |
| Cell dissociation | Organoids dissociated with Accutase; rods left at the bottom | 20 min incubation |
| Filtration | Cells passed through a 40 µm strainer | — |
| Resuspension | Cells in PBS + 0.1% BSA | — |
| Library preparation & sequencing | 10x Genomics platform | — |
| Initial processing | Cell Ranger used to generate count matrix | — |
| Seurat object creation | Count matrix converted into Seurat object | — |
| Filtering: gene counts | Remove cells with too few or too many genes | <2,000 or >10,000 genes |
| Filtering: molecule counts | Remove cells with extreme total molecule counts | <5,000 or >100,000 molecules |
| Filtering: mitochondrial reads | Remove cells with high mitochondrial content | >10% mitochondrial reads |
| Doublet removal | Identify and remove doublets | *scDblFinder* package |
| Normalization | Normalize gene expression | NormalizeData function, Seurat |
| Highly variable gene selection | Identify genes for downstream analysis | Top 2,000 genes |
| Scaling | Scale datasets to reduce technical variation | — |
| Linear dimensionality reduction | Principal component analysis (PCA) | Up to 30 PCs used for clustering |
| Clustering | Identify cell clusters | Resolution = 0.5 |
| Non-linear dimensionality reduction | UMAP embedding | — |
| Cluster annotation | Manual annotation using canonical marker genes | See Table 7.1 |

Gene List For the annotation of hepatic lineages:
| Cell types | Description | Gene list used for classification |
| --- | --- | --- |
| **HE** | Hepatic endoderm | FOXA2, TBX3, KRT8, KRT18, EPCAM |
| **efHB** | Early fetal Hepatoblasts | EFNB2, TTC6, SERPINA1 |
| **fHB** | Fetal hepatoblast | ONECUT2, HHEX, PROX1, NPW |
| **pHB1** | Proliferative Hepatoblast (in the process of proliferation) | UBE2C, CDC20, CENPF, SOX17 |
| **pHB2** | Proliferative Hepatoblasts 2 (exit mitotic cell cycle) | CLSPN, HIST1H1A, MCM10 |
| **miHB** | Migratory Hepatoblasts | GATA4, CDH2, FN1, GPC3 |
| **fHP** | Foetal hepatocytes | ALB, SERPINA6, H19, AFP |
| **HP1** | Immature Hepatocytes | ALDH6A1, HNF4A, AFP, APOA1, APOM, PEBP1, B2M, FTL |
| **HP2** | Mature hepatocytes | APOA4,SLC10A1,GSTA1,ABCC2,APOC3,AHSG,HPX,RBP, AMN |
| **BEC** | Biliary Epithelial cells | KRT7, KRT19,EPCAM,SOX9,CLDN4,CFTR,CDH1 |
| **pHSC** | Proliferative Hepatic stellate cells | CDK1, CENPF,RRM2, NUSAP1, COL1A1, COL3A1, COL6A3, ITGA8, MMP2, ZEB2, EPHA3, MFAP4, TNNI1 |
| **aHSC** | activated Hepatic stellate cells (Collagen production) | COL1A1, COL3A1, COL6A3, ITGA8, MMP2, ZEB2, EPHA, MFAP4,TNNI1 |
| **EC** | Endothelial cells | PECAM, OTI3, KDR, GNG11, CDH5, NOTCH1 |

