## Supplementary material for "Bioscaffold guidance drives liver periportal area tubulogenesis in hIPSC organoids": supplemetary figures

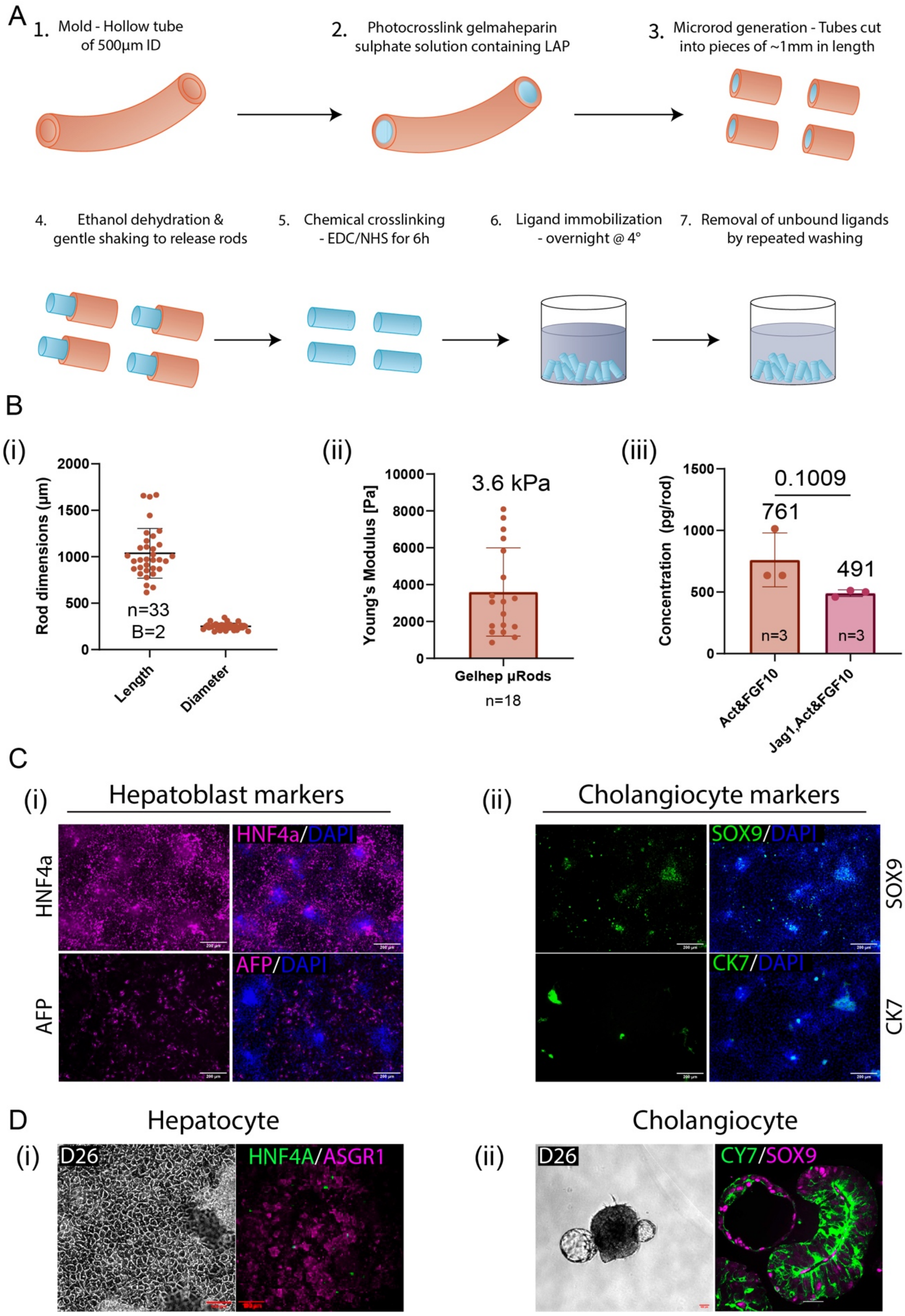


**Supplementary Figure 1:**

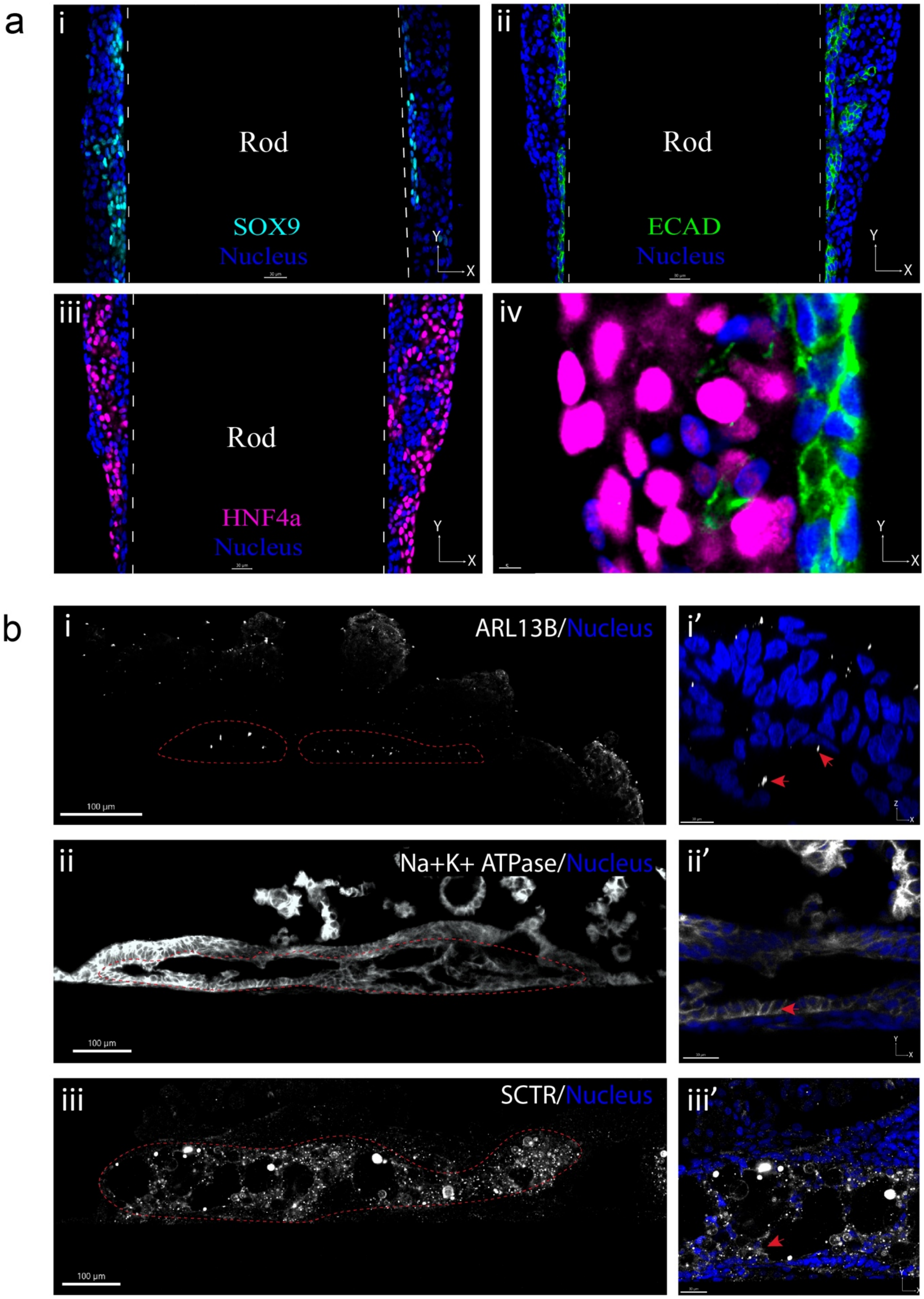


**Supplementary Figure 2:**

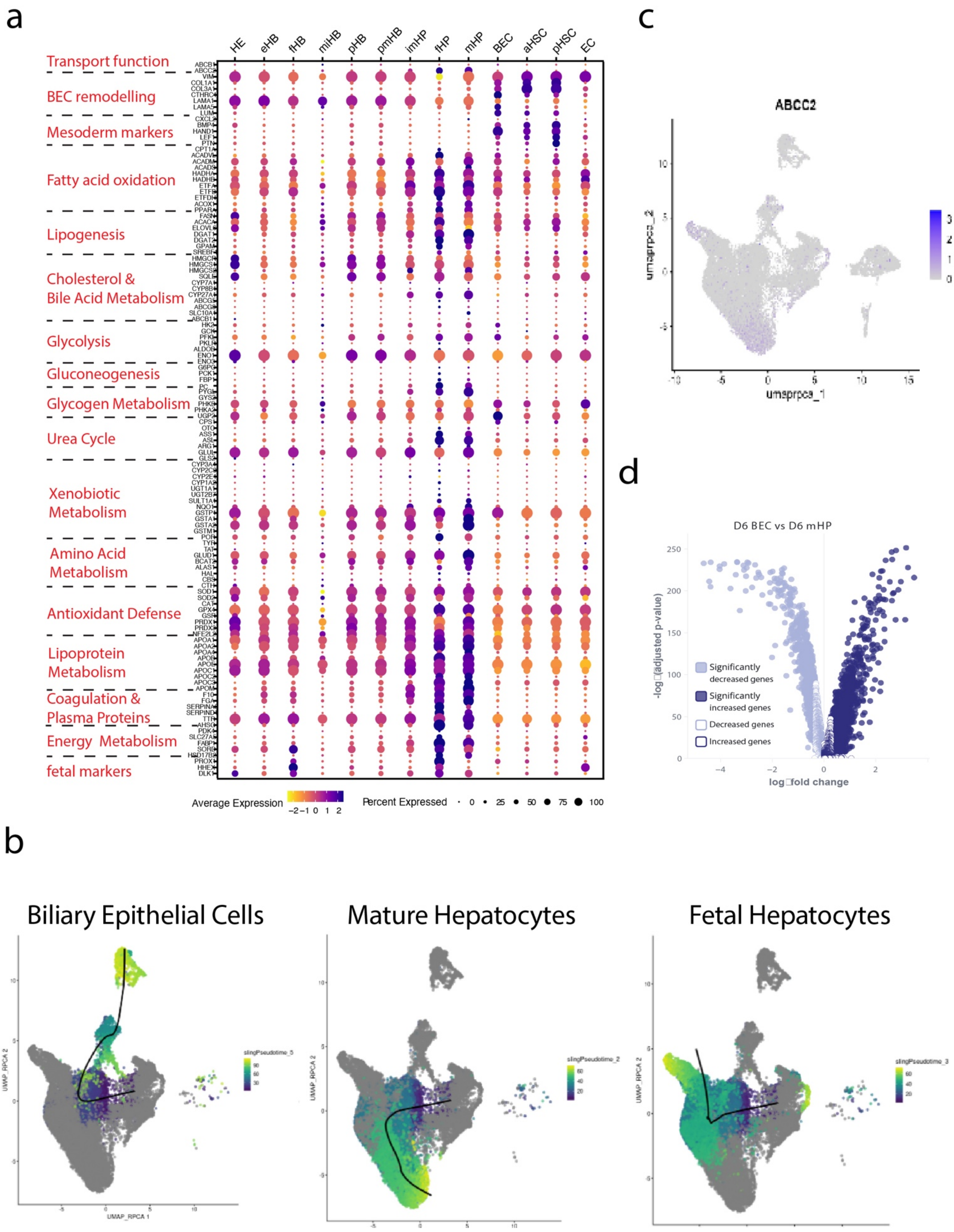


**Supplementary Figure 3:**

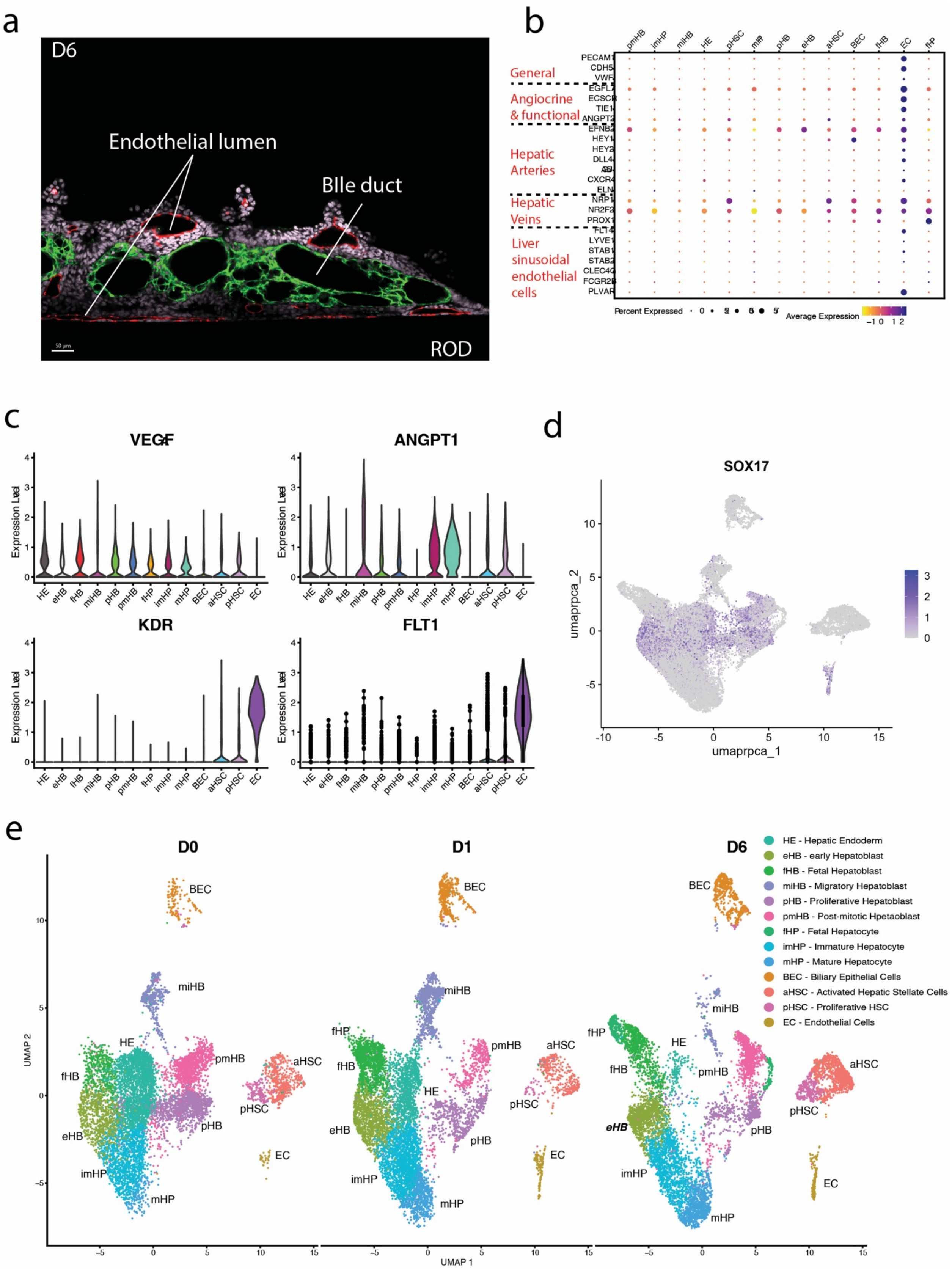


**Supplementary Figure 4:**

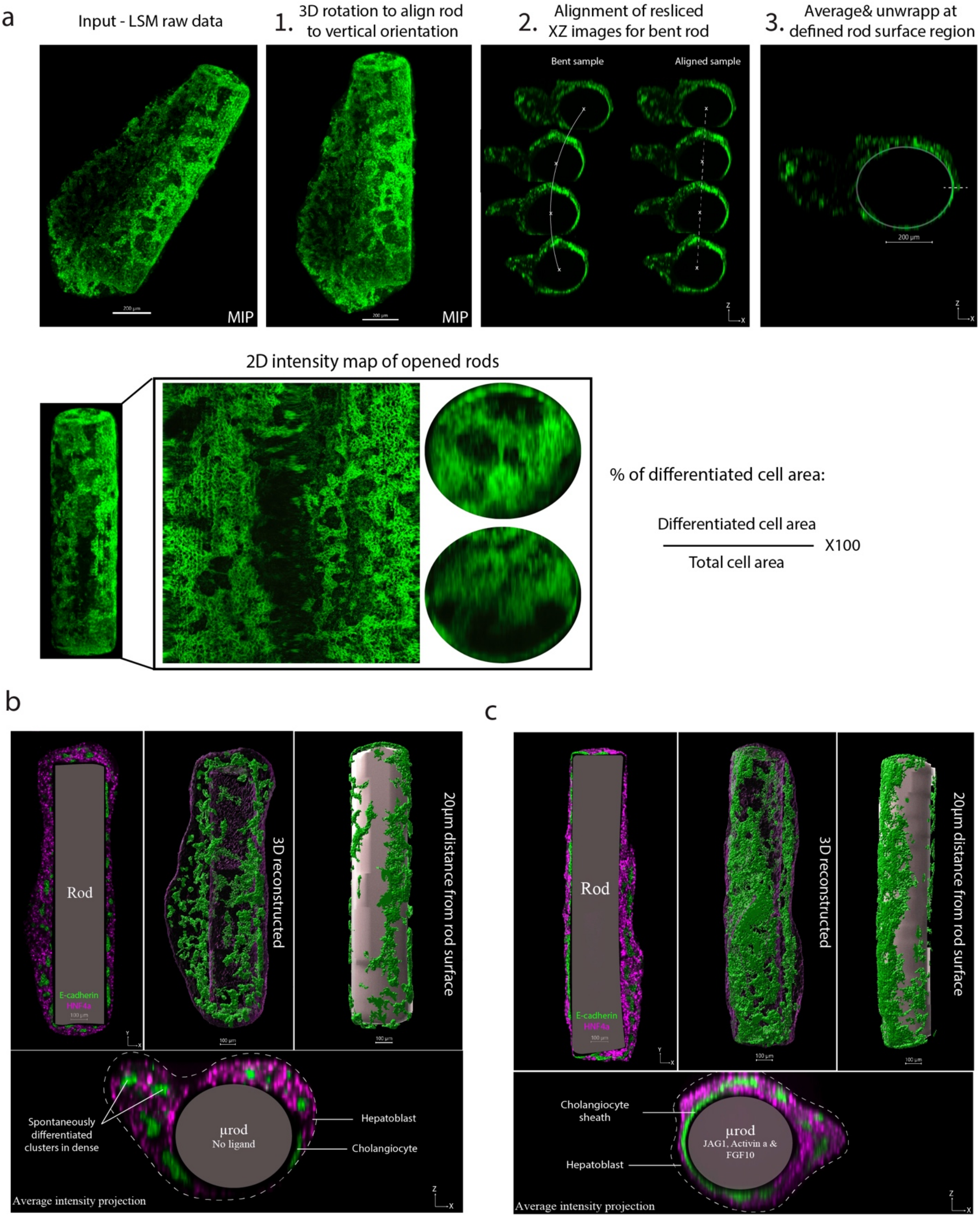


**Supplementary Figure 5:**

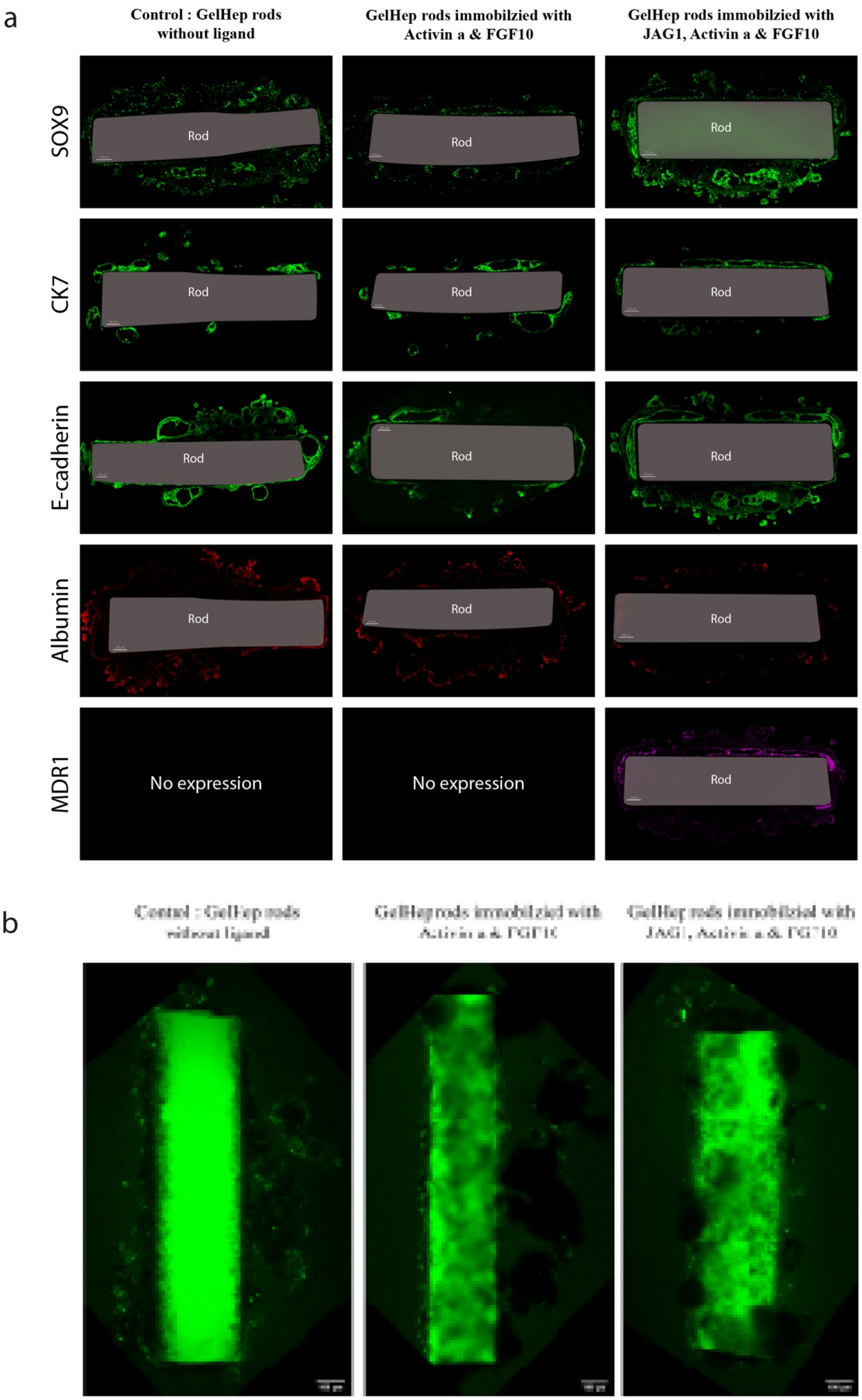


**Supplementary Figure 6**:

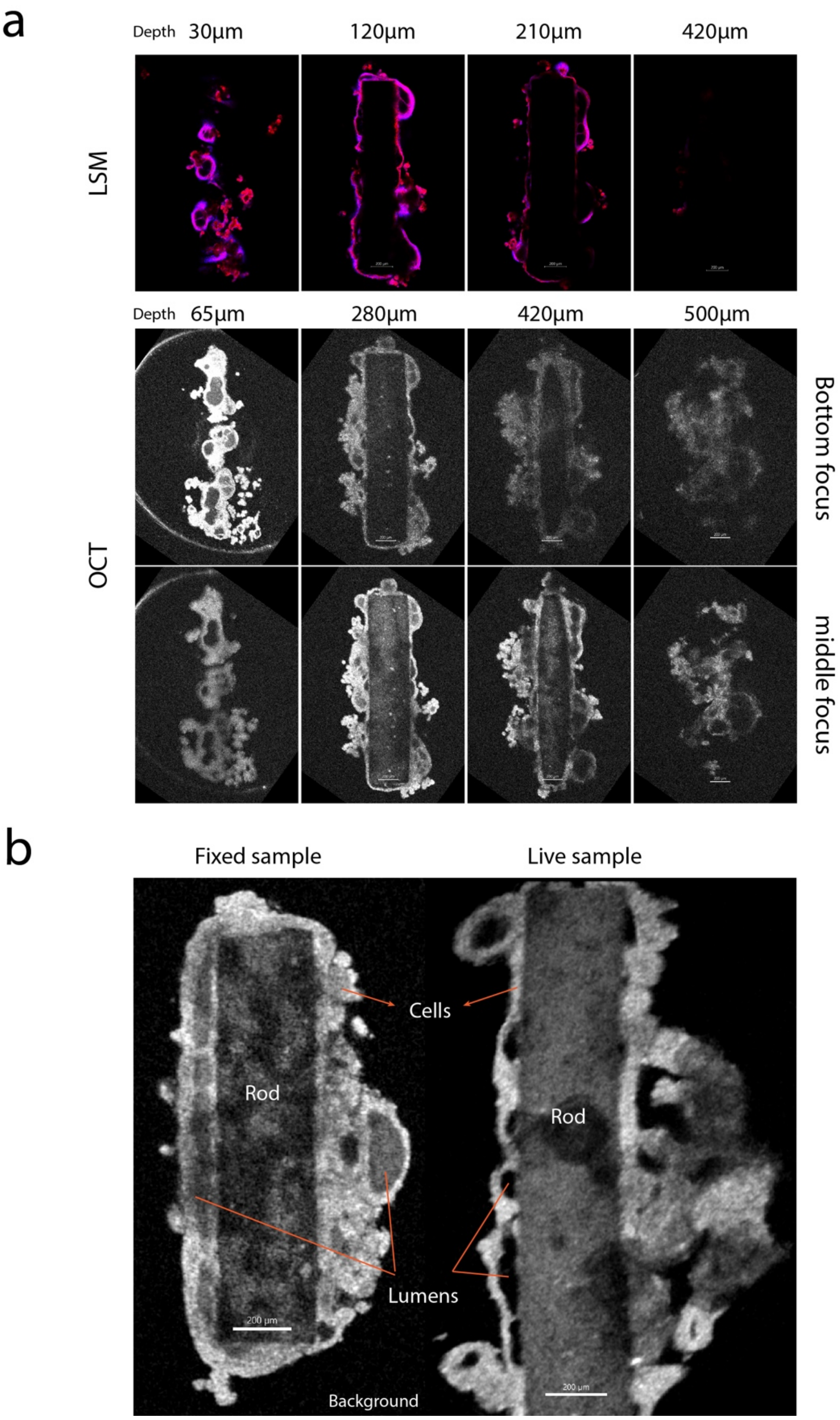


**Supplementary Figure 7:**

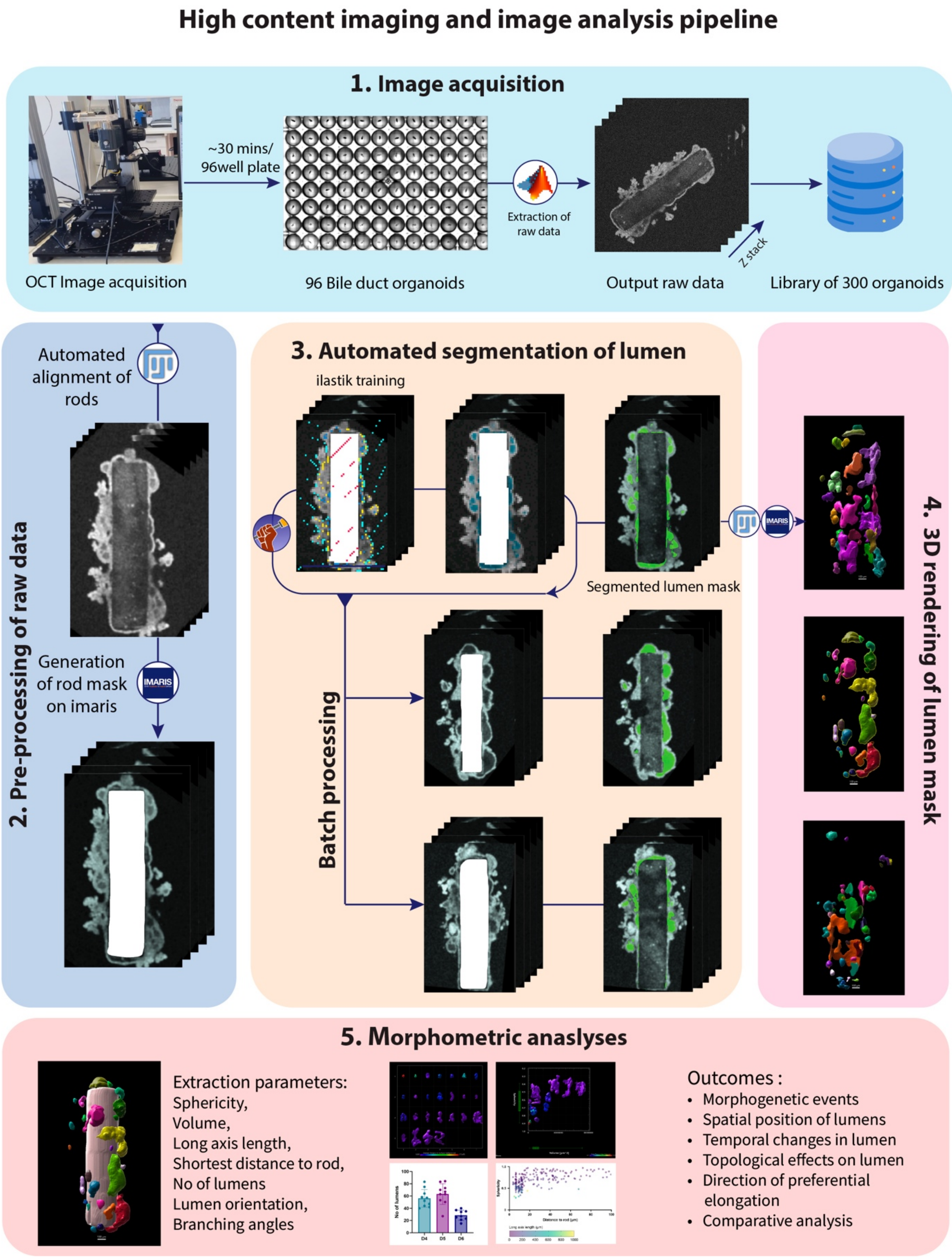


**Supplementary Figure 8:**

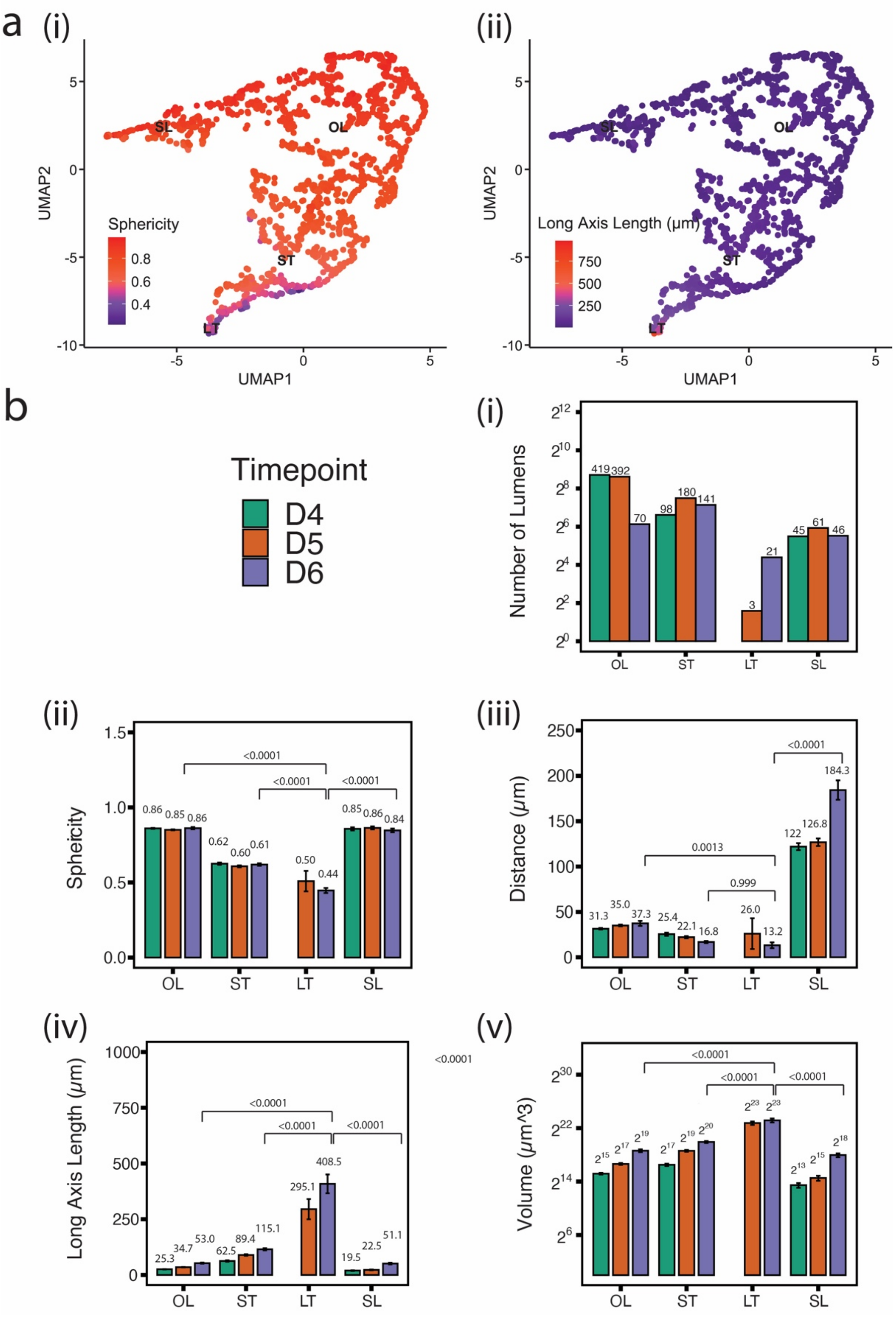


**Supplementary Figure 9:**

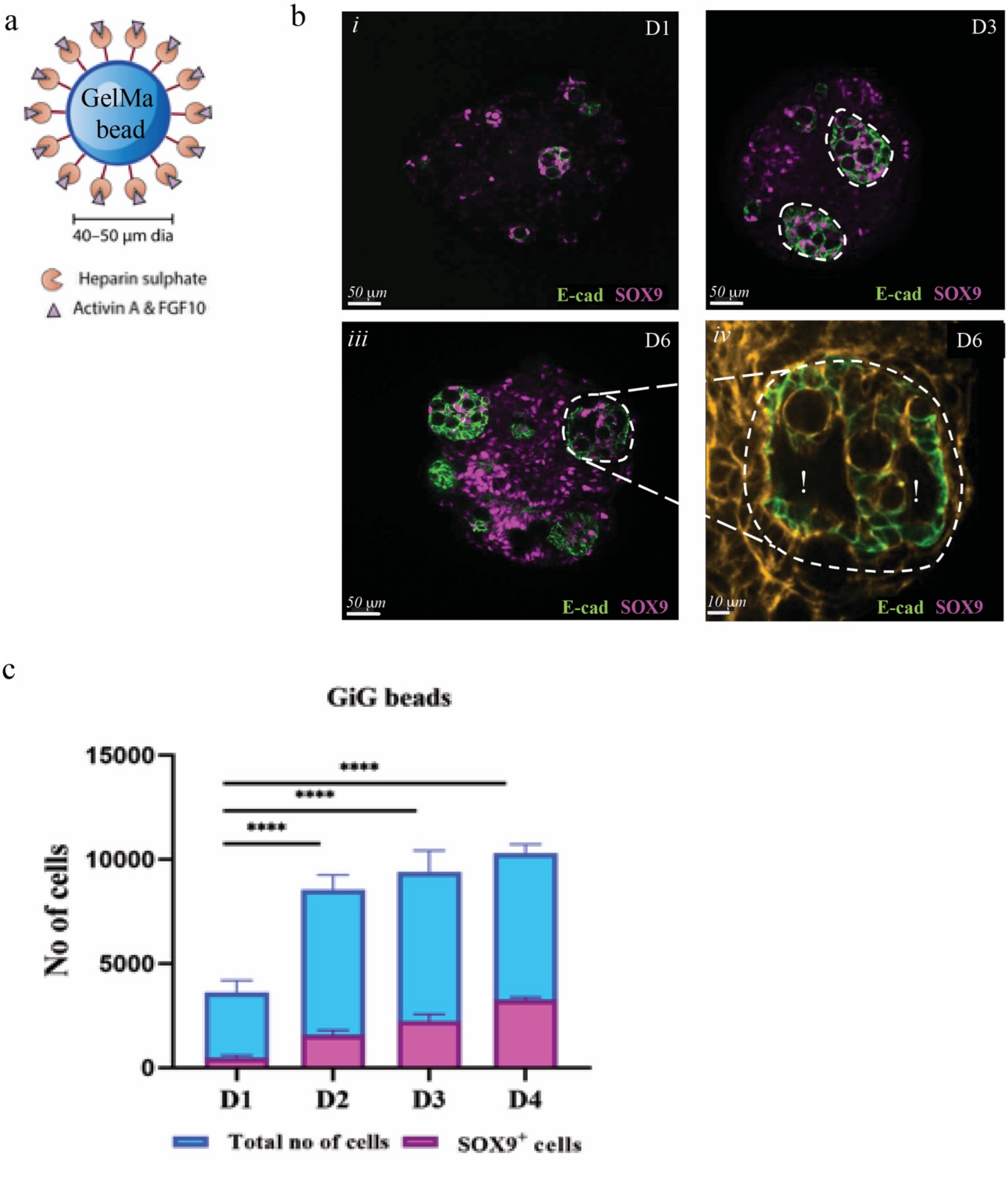


**Supplementary Figure 10:**

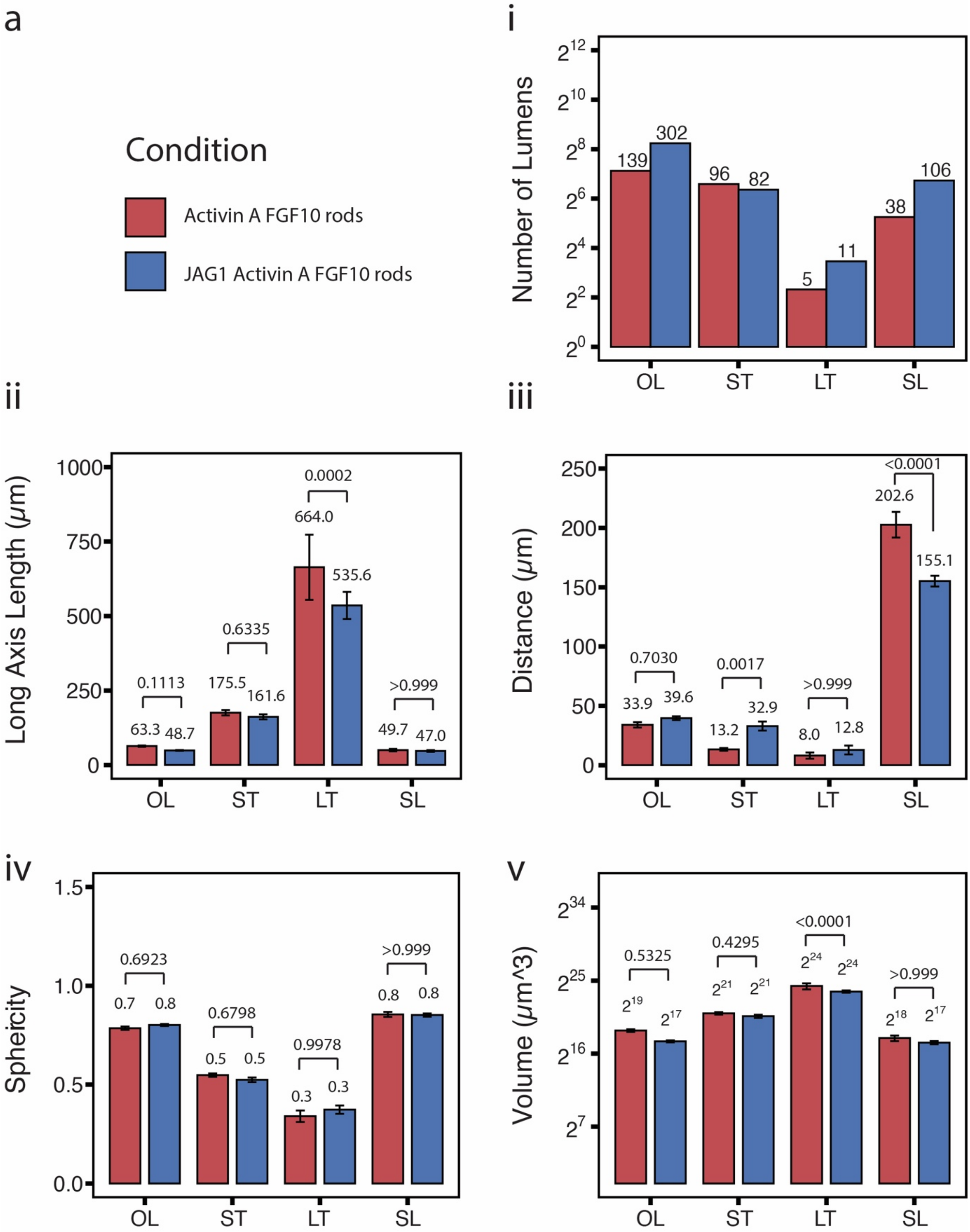


**Supplementary Figure 11:**
